# Dynamic protein assembly and architecture of the large solitary membraneless organelle during germline development in the wasp *Nasonia vitripennis*

**DOI:** 10.1101/2023.12.13.571440

**Authors:** Kabita Kharel, Samuel J. Tindell, Allie Kemph, Ryan Schmidtke, Emma Alexander, Jeremy A. Lynch, Alexey L. Arkov

## Abstract

Germ cells in different animals assemble characteristic membraneless organelles referred to as germ granules, which contain RNA and proteins required for germline development. Typically, the germ granules are small spherical or amorphous cytoplasmic granules and often, they assemble around membrane-bound organelles such as nuclei, mitochondria and endoplasmic reticulum. In particular, in egg chambers of the fruit fly *Drosophila*, nurse cells assemble perinuclear granules, referred to as nuage, along with multiple small germ granules formed at the posterior pole of the oocyte (polar granules). Nuage is assembled in a very similar way in the wasp *Nasonia vitripennis*, despite the long evolutionary distance from *Drosophila.* In contrast, *Nasonia* forms a very different single germ granule, called the oosome, at the posterior, which is about 40 times larger than a homologous *Drosophila* polar granule. Here, using molecular and super-resolution imaging approaches, we provide insights into protein assembly and architecture of the oosome during germline development. Interestingly, unlike the fly, the wasp utilizes alternatively spliced RNA-helicase Vasa isoforms during germline development and oosome formation. The isoforms differ by an unstructured region, containing repeats of phenylalanine and glycine, that is similar to functional domains characteristic of nucleoporins. In addition, while other conserved components of germ granules, such as Oskar, Aubergine and Tudor proteins are recruited to the oosome, these polypeptides show a distinct and specific localization within the oosome. Of particular note, Tudor protein forms a shell encapsulating the oosome, while small Oskar/Vasa/Aubergine granules occur inside the oosome core. Also, in surprising contrast to *Drosophila* egg chambers, we found that a subset of the wasp nurse cells located in anterior show dramatic DNA damage and assemble higher levels of nuage than their posterior counterparts. The characteristics of two distinct nurse cell populations suggest a mechanistic link between the higher amounts of nuage assembled in anterior nurse cells and their need to silence transposable elements in the presence of double-strand DNA breaks. Our results point to the high degree of plasticity in the assembly of membraneless organelles, which adapt to specific developmental needs of different organisms, and suggest that novel molecular features of conserved proteins result in the unique architecture of the oosome in the wasp.

## Introduction

Membraneless RNA-protein granules are assembled in all living systems and cell types and may represent one of the earliest levels of biological organization which has predated the cellular life^1^. Typically, the membraneless organelles are numerous and small structures (ranging from a few hundred nanometers to a few micrometers in diameter) and often their assembly process follows the principles of phase separation which occurs during formation of macromolecular condensates^2–5^.

In germ cells of different animals, specific cytoplasmic membraneless granules form from RNA and proteins required for germ cell specification^6–12^. These germ granules often associate with other organelles including the nuclei, mitochondria and endoplasmic reticulum^13,14^. In particular, in the fruit fly *Drosophila*, during oogenesis, germ granule nuage is assembled at the cytoplasmic side of the nuclear envelope of nurse cells in developing egg chambers. Also, *Drosophila* oocyte assembles specialized cytoplasm - germ plasm - at the posterior pole, which contains multiple small germ granules of approximately 500 nm in diameter referred to as polar granules. The germ plasm and polar granules remain positioned at the posterior pole during early embryogenesis before they are segregated into primordial germ cells during their formation at the posterior pole^15^.

In stark contrast to multiple polar granules in *Drosophila*, the small wasp *Nasonia vitripennis* assembles a large solitary membraneless organelle referred to as the oosome, which is about 40 times larger than the typical polar granule^16^. Initially formed in the posterior of the oocyte, the oosome migrates within the cytoplasm of the early embryos before returning to the posterior pole where it breaks down to segregate into primordial germ cells^16^.

Here we show that, regardless of the dramatic difference in size, both polar granules and the oosome incorporate several conserved proteins including Oskar (Osk)^17–20^, Vasa (Vas)^21^, Aubergine (Aub)^22,23^, and Tudor (Tud)^24–26^. However, our molecular analysis of these conserved proteins uncovered unexpected distinctions in these polypeptides between the fly and the wasp. In particular, we provide evidence for the presence of a novel N-terminal segment of *Nasonia* Oskar (Nv-Osk) adjacent to the conserved LOTUS domain that is absent in other insect Osk proteins. In addition, we found a novel Vas splice isoform in the wasp. This new isoform is the most prevalent form of *Nasonia* Vasa (Nv-Vas) found in the early embryo. Interestingly, we show that this Nv-Vas isoform contains an intrinsically disordered Phe-Gly (FG)-rich segment, which, while not present in the fly Vas, it is found in nucleoporin proteins of the nuclear pore complex^27–29^ and Vas homologs of the worm *C. elegans*^30–32^ where it plays an important role in regulating the properties of phase separated condensates.

Furthermore, we systematically characterized the subcellular localization of this Nv-Vas isoform and other germ granule protein components during germline development. Unexpectedly, we found that *Nasonia* Tudor (Nv-Tud) protein is highly enriched at the periphery of the oosome, and forms a fibrous shell which envelops the oosome core. Interestingly, Nv-Osk, Nv-Vas and *Nasonia* Aubergine (Nv-Aub) are assembled in the small spherical granules in the oosome core, which, while similar in size to polar granules in *Drosophila* germ plasm, show distinct morphology and mechanisms of assembly. We further show that the Nv-Tud shell breaks down as the embryonic nuclei begin to interact with the oosome at the onset of the primordial germ cell formation. Our data suggest that the Nv-Tud shell provides mechanical stability to contain a less dense oosome core during its migration in the early embryo. In addition, we show that contrary to *Drosophila* oogenesis, *Nasonia* egg chambers have distinct subset of nurse cells in anterior that show evidence of double-strand DNA breaks and assemble higher amounts of perinuclear nuage than their posterior counterparts, indicating a higher demand for anterior nurse cells to silence transposable elements in the presence of DNA damage. Overall we find that a combination of novel protein features and redeployment of conserved elements have combined in the evolution of the highly unusual subcellular organelle in the wasp.

## Results

### *Nasonia* germ cells expresses two isoforms of Vasa, which differ by the inclusion of an FG-rich region and show differential expression in the ovary and early embryo

To begin characterization of proteome of the oosome, we generated antibodies for the wasp homolog of the broadly conserved germ granule marker and ATP-dependent RNA helicase Vas (Nv-Vas). Interestingly, rabbit anti-Nv-Vas antibody detected two proteins at ∼80-100 kD range (Fig. 1a). Both proteins were reduced in an *Nv*-*vas* RNAi experiment (Fig. 1b) indicating that those two proteins correspond to two Nv-Vas isoforms. Furthermore, we were able to IP both proteins with this antibody, demonstrating that the detection of an extra anti-Nv-Vas-reactive band is not caused by generating an anti-Nv-Vas reactive epitope in an unrelated protein due to denaturing conditions of a western-blot experiment (Extended Data Fig. 1a). Subsequently, we were able to confirm Nv-Vas identity of both IP-ed proteins with mass spectrometry.

**Fig. 1.**
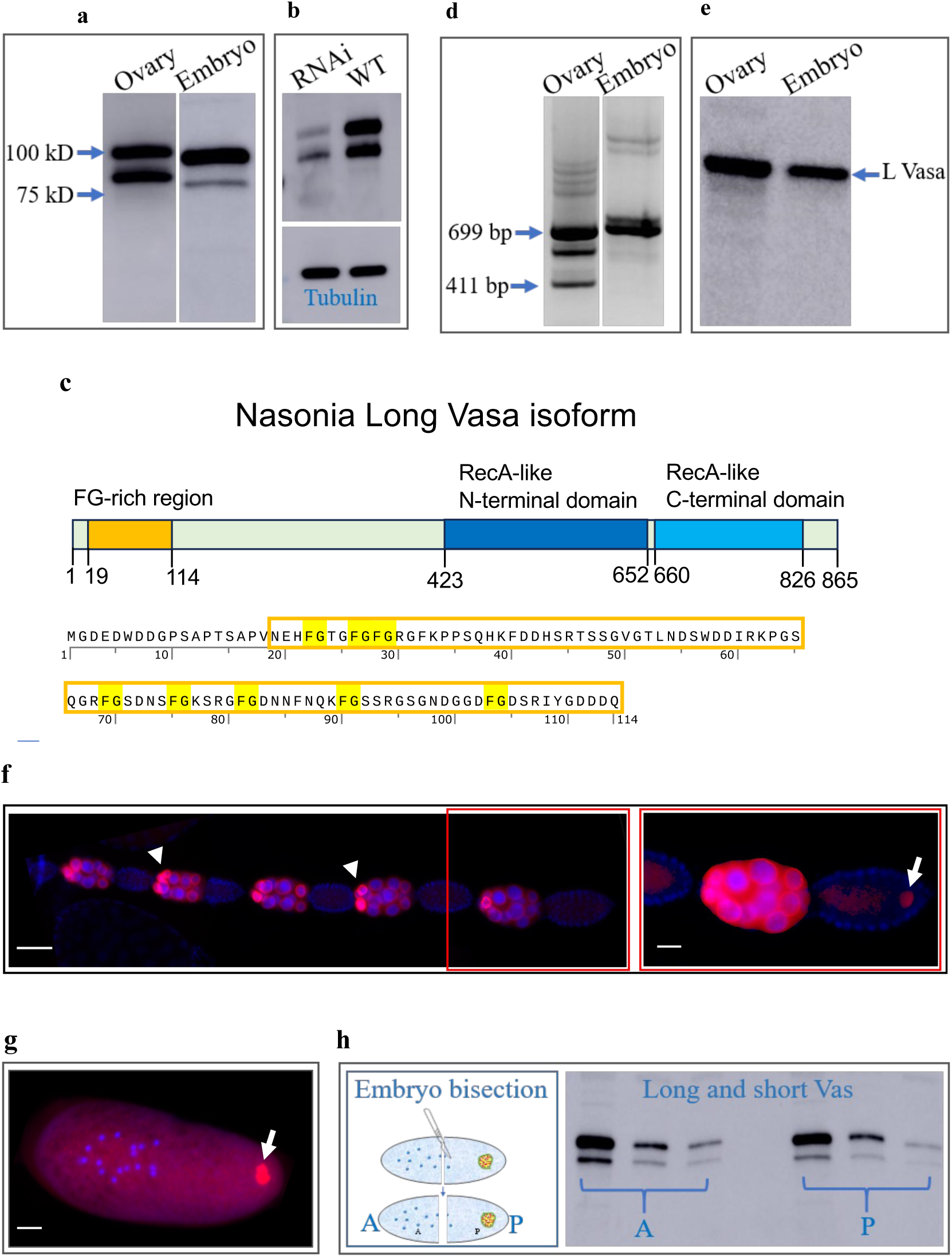
*Nasonia* expresses two isoforms of Vasa protein, which differ by the Phe-Gly-rich region and show preferential expression in anterior nurse cells. **a, b**, Two isoforms of Nv-Vas in *Nasonia*. **a**, Detection of two isoforms in *Nasonia* ovary and embryos using rabbit anti-Nv-Vas antibody in a western blot experiment. **b**, Reduction of Nv-Vas is observed in *Nv*-*vas* RNAi ovarian samples compared to wild-type control samples (top), equal loading was confirmed with anti-β-Tubulin antibody (bottom). **c**, Top: a diagram of a Long isoform of Nv-Vas protein (UniProt ID A0A7M7G4Z2) indicating the correct locations of N-terminal Phe-Gly (FG)- rich region, which is lacking in the Short Nv-Vas isoform. Also, the location of the conserved RecA-like domains required for the ATP-dependent RNA helicase activity are shown based on the structure of *Drosophila* Vas^59^ and alignment of multiple Vas proteins^60^. Bottom: Amino acid sequence of N-terminal region of Long Nv-Vas, which includes this isoform-specific FG-rich region (amino acids 19-114), indicated with box. **d**, RT-PCR experiment using RNA from *Nasonia* ovarian and embryonic extract confirms the presence of RNA for the two Nv-Vas isoforms. The Short *Nv-vas* isoform RNA (corresponds to 411 bp RT-PCR product) is detected in the ovary but it is not observed in the embryo. The Long *Nv-vas* isoform RNA (corresponds to 699 bp RT-PCR product) is detected in ovary and it is the only *Nv*-*vas* RNA persisted in early embryos. Identity of both *Nv-vas* isoforms was confirmed by direct sequencing of the corresponding RT-PCR fragments. **e**, Antibody raised against the FG-rich region (amino acids 19-114), recognizes only the Long Nv-Vas isoform in both ovary and embryo in western blot experiments. **f**, Left: Developing egg chambers in a single ovariole stained with anti-Long Nv-Vas antibody (red) and nuclear stain DAPI (blue). The Nv-Vas is strongly enriched in anterior nurse cells (arrowheads). Scale bar, 50 μm. Right: A higher magnification image of the last egg chamber shown in the left panel is indicated in red box. The cortically anchored oosome at the oocyte’s posterior labeled with anti-Long Nv-Vas becomes first visible at early stage 3 of oogenesis as indicated with arrow. This image was obtained with high signal intensity to show the oosome, which has significantly less Nv-Vas at this early stage compared to the nurse cells. Scale bar, 20 μm. **g**, *Nasonia* preblastoderm embryo showing oosome in the posterior part of the embryo (arrow). Oosome is stained with anti-Long Nv-Vas antibody (red). DAPI stains nuclei (blue). Scale bar is 30 μm. In **f**, **g**, anterior is to the left. **h**, Left: a schematic illustrating the bisection of the embryo to produce anterior (A) and posterior (P) halves. Right: western blot analysis shows the detection of Long and Short Nv-Vas isoforms in both the anterior (oosome lacking) and posterior (with the oosome) sections of the embryos. Three different protein amounts were used for each section and show equal expression of Nv-Vas in both anterior and posterior regions.

Next, using the rabbit anti-Nv-Vas antibody, we compared Nv-Vas expression in embryos versus ovaries and we found that contrary to the ovary, the Short Nv-Vas isoform was substantially decreased in the embryos (Fig. 1a).

Since, in *Drosophila*, Vas is known to be phosphorylated^33,34^, the expression of two isoforms could be explained by a phosphorylation of Nv-Vas in *Nasonia*. However, the treatment of ovarian lysate with CIP enzyme, which removes phosphates, failed to eliminate the presence of either of the Nv-Vas bands (Extended Data Fig. 1b) suggesting that phosphorylation is not the cause of an extra Nv-Vas isoform in *Nasonia*.

Computational analysis predicted two alternatively spliced *Nv*-*vas* mRNAs, that should result in 92.3 kD and 82 kD proteins. Interestingly, the predicted Long Nv-Vas isoform has an 96 amino acid insertion, which contains multiple repeats of phenylalanine and glycine (FG repeats) (Fig. 1c and Extended Data Fig. 1c), which are commonly detected in nucleoporins^27–29^ and also, in the worm *C. elegans* Vas homologs^30–32^ but not in *Drosophila* Vas. Therefore, we tested whether these two predicted isoforms are expressed in the germline. RT-PCR experiments showed that ovaries express RNA for both alternatively spliced isoforms (Fig. 1d). However, in the early embryonic extracts only the Long *Nv-vas* RNA isoform was detected (Fig. 1d). This was consistent with Long Nv-Vas being the major isoform in early embryos (Fig. 1a).

To further determine whether these two isoforms are encoded by the predicted alternatively spliced *Nv-vas* RNAs and characterize Nv-Vas in more detail, we generated a guinea pig Long Nv-Vas-specific antibody using the 96 amino acid protein fragment that is missing in the Short Nv-Vas isoform, as an antigen. Accordingly, Fig. 1e shows that the Nv-Vas Long isoform-specific antibody recognized a single protein, which is identical to the Long Vas isoform that was detected with rabbit anti-Nv-Vas antibody (Fig. 1a).

To determine the distribution of Long Nv-Vas isoform in ovary and early embryos, we immunostained these tissues with the Long Nv-Vas-specific antibody (Fig. 1f, g). Interestingly, this isoform is enriched in a small population of anterior nurse cells early during oogenesis and later begins to localize to the oosome at the oocyte’s posterior in early stage 3 of oogenesis (*Nasonia* oogenesis staging was done as described previously^35,36^) (Fig. 1f right panel). The distribution of Long Vas isoform shown here and that of both Long and Short isoforms together detected by our new rabbit antibody and previously generated antibody against a peptide common to both isoforms, in the ovary are very similar^18^, indicating that both isoforms are distributed together during oogenesis.

In the early (preblastoderm) embryos, the Long Nv-Vas isoform is incorporated in the oosome (Fig. 1g). In addition, we detected Nv-Vas outside the oosome, distributed in the embryo’s cytoplasm (Fig. 1g). To further confirm the Nv-Vas distribution pattern in the early embryo, we cut the embryos in half and compared the Nv-Vas content in anterior half (lacks the oosome) and oosome-containing posterior half. Mass spectrometry and western-blot analysis identified Nv-Vas in both anterior and posterior halves (Fig. 1h), indicating that, in addition to the strong localization of Nv-Vas to the oosome, this protein is expressed outside the oosome.

The presence of FG-rich region in a major Nv-Vas isoform in the embryos, which is absent in *Drosophila* Vas, raised a possibility that, similar to nucleoporins, the Nv-Vas FG-rich region is intrinsically disordered. This may contribute to the formation of size-exclusion mesh-like assemblies by engaging aromatic groups of phenylalanine residues in intermolecular *π*-*π* interactions similar to those of the nuclear pore complex and *C. elegans* Vas homologs.

Therefore, to test the structure of this region, we purified an Nv-Vas tagless FG-rich region under native conditions and probed its structure with Circular Dichroism (CD) spectroscopy (Extended Data Fig. 2). Interestingly, CD spectrum of this region was typical to the spectra of unfolded proteins and, specifically, closely resembles intrinsically disordered FG-repeat regions of nucleoporins^27,28^ with large ellipticity minimum close to 200 nm and low ellipticity around 230 nm region (Extended Data Fig. 2c). Therefore, these data provide experimental evidence that the alternatively spliced Vas isoforms in *Nasonia* differ by an intrinsically disordered FG-rich region which may be involved in the formation of large Vas assemblies in *Nasonia* germ cells and early embryos.

### *Nasonia* Oskar protein contains novel domain and it forms small granules in the core matrix of the oosome and *Drosophila* S2 cells

In *Drosophila*, Vas is recruited by Osk to the germ plasm^37,38^. Therefore, we generated an Nv-Osk-specific antibody to investigate the relationship between Vas and Osk in *Nasonia* and to gain detailed insights into the molecular assembly pathway of the oosome. Unexpectedly, instead of 45 kD Nv-Osk protein, predicted earlier, we detected a longer (∼51 kD) protein with the antibody (Fig. 2a). Furthermore, this 51 kD protein was reduced in *Nv-osk* RNAi experiments validating the specificity of the antibody and confirming that the 51 kD protein is Nv-Osk (Extended Data Fig. 2a). In addition, in different experiments, we were able to IP the 51 kD Nv-Osk from both ovarian and embryonic extracts using our antibody under native conditions (Extended Data Fig 2b). Therefore, we asked whether there are some additional uncharacterized exon regions at the 5′ and 3′ ends of *Nv-osk* mRNA which could be translated to account for an extra ∼6 kD extension. The *Nv-osk* mRNA 3’ RACE mapping was consistent with the previously identified 3′-end of *Nv-osk* mRNA indicating that Nv-Osk is not extended at its C-terminus. However, 5′-end mapping revealed that *Nv-osk* RNA starts at more than 800 bps upstream of previously predicted *Nv-osk* transcription start site (Extended Data Fig. 2c). The extended 5′ UTR codes for previously unrecognized 57 additional amino acids (6.6 kD), which we refer to as Nv-Osk N-terminal domain (NTD), immediately upstream of the LOTUS domain of Osk (Fig. 2b). To confirm that the NTD is responsible for the observed higher molecular weight of *Nasonia* Osk, we expressed new full-length coding sequence of *Nv*-*osk* (which included NTD) in bacterial in vitro translation system, and observed identical migration of Osk generated in vitro to the 51 kD protein detected in the ovary and embryos (Extended Data Fig. 3d). Interestingly, in some western-blot experiments with anti-Nv-Osk antibody using both ovarian and embryonic lysates, in addition to major 51 kD band, a less intense ∼100 kD band is detected, suggesting the formation of Osk dimers (Extended Data Fig. 3e) consistent with previous finding that LOTUS domain of Nv-Osk dimerizes^17^.

**Fig. 2.**
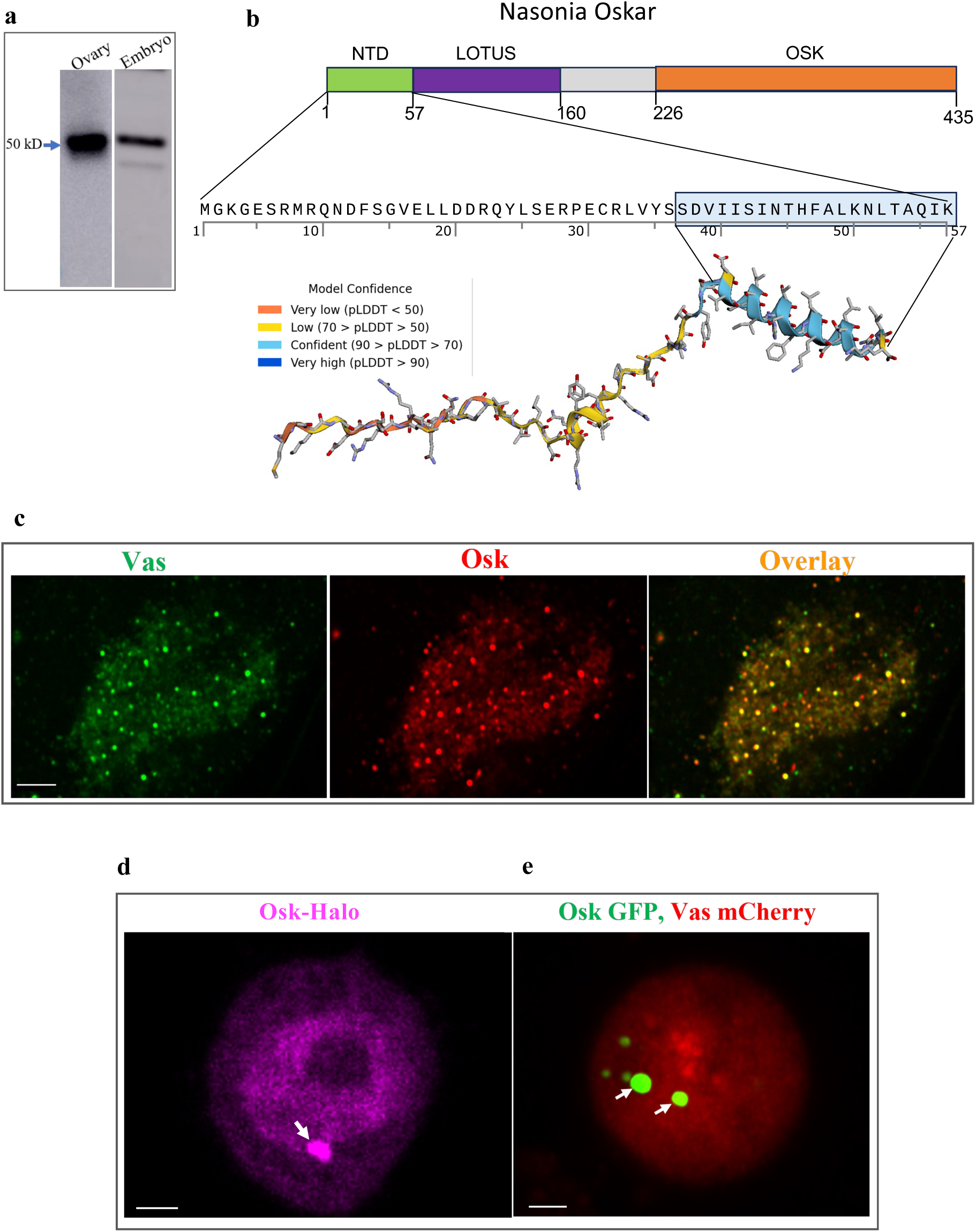
*Nasonia* Oskar contains novel N-terminal domain and is assembled into small granules in the matrix of the oosome and *Drosophila* S2 cells. **a**, anti-Nv-Osk antibody reacts with protein which is ∼6 kD longer than expected in both *Nasonia* ovary and early embryos. **b**, Molecular analysis identified a long 5′ UTR in *Nv-osk* which codes for novel N-terminal domain (NTD) (Extended Data Fig. 3 c, d). Amino acid positions of LOTUS and OSKAR domains of Nv-Osk are indicated. Sequence of the NTD and its AlphaFold-predicted structural model with model confidence values are shown. The model predicts a long α−helix adjacent to the LOTUS domain (highlighted in blue). **c**, Super-resolution confocal microscopy image of the oosome in preblastoderm embryos demonstrating that Nv-Vas, stained with rabbit anti-Nv-Vas antibody (green channel) and Nv-Osk, detected with guinea pig anti-Nv-Osk antibody (red channel) are co-assembled in the small granules inside the oosome matrix. (**d**, **e**), Nv-Osk is assembled in the granules in *Drosophila* S2 cells (indicated with arrows) irrespective of a fluorescent tag (Halo tag in **d** and GFP in **e**) and of the presence of Nv-Vas proteins (**d**, without Nv-Vas isoforms; **e**, in the presence of both Short and Long Nv-Vas isoforms tagged with mCherry). Scale bar in **c** is 3 μm and in (**d**, **e**) it is 2 μm.

The Nv-Osk NTD appears to be a novel region since we could not find similar sequences in genomes of different organisms. Interestingly, AlphaFold^39^ predicts a long α−helix in the NTD adjacent to the conserved LOTUS domain of Osk (Fig. 2b), which may be a novel structural extension of the *Nasonia* Osk LOTUS domain (Fig. 2b).

Next, we employed super-resolution microscopy to analyze the pattern of Nv-Osk and Nv-Vas in preblastoderm embryos in detail. In particular, simultaneous detection of both proteins revealed that they are incorporated into small (∼400 nm) spherical granules inside the oosome, along with a fibrous network of protein that extends through the bulk of the oosome (Fig. 2c).

In order to determine whether Nv-Osk and Nv-Vas require the molecular context of the oosome to form the granules, we tested whether these proteins can form the granules when expressed in a heterologous system - *Drosophila* somatic Schneider S2 cells. Fig. 2 shows that Nv-Osk was able to assemble into spherical granules in S2 cells, when expressed alone (Fig. 2d) or in the presence of Nv-Vas protein isoforms (Fig. 2e). These granules resemble Nv-Osk granules formed in the embryonic oosome matrix. However, contrary to similar experiments for *Drosophila* Osk and Vas^40^, Nv-Vas proteins were not enriched in Nv-Osk granules indicating that, contrary to *Drosophila*, Nv-Osk does not directly recruit or associate with Nv-Vas in *Nasonia* (Fig. 2e). Furthermore, we could not detect Nv-Osk in Nv-Vas immunoprecipitates (Extended Data Fig. 3f) providing further support for the lack of strong interaction between Nv-Osk and Nv-Vas in the wasp. Interestingly, consistent with our experiments suggesting the lack of direct interaction between Nv-Vas and Nv-Osk, we noticed that some amino acids in *Drosophila* Vas, important for interaction with Osk or for Vas localization to germ granules, have not been conserved in Nv-Vas. Specifically, Phe508 in *Drosophila* Vas, shown to be important for interaction with Osk^38^, is changed to Leu in the corresponding position 711 of Nv-Vas. In addition, Gln527 in *Drosophila* Vas is required for Vas recruitment to germ granules in *Drosophila*^41^, and it corresponds to Glu at position 730 of Nv-Vas (Extended Data Fig. 1c).

In *Drosophila*, Vas expression in S2 cells induces the formation of granules by itself^40^. However, we found that Nv-Vas long and short isoforms, when either expressed separately or co-expressed in S2 cells, fail to form granules (Fig. 2e). Overall, our data suggest that while Nv-Osk has the intrinsic ability to condense into spherical granules, Nv-Vas may not interact with Nv-Osk directly and it may require other oosome components to assemble with Osk in the oosome. Thus, in contrast to *Drosophila*, *Nasonia* may exhibit a distinct mechanism during the assembly of these proteins into granules.

### Distinct populations of nurse cells in *Nasonia* egg chambers are different from *Drosophila* and show different amounts of perinuclear nuage and double-strand DNA breaks

The striking enrichment of Nv-Vas in a small subset of anterior nurse cells of the egg chamber before its assembly into the oosome at the oocyte’s posterior is different from *Drosophila* egg chambers, which show equal distribution of Vas in all 15 nurse cells of an egg chamber in the perinuclear nuage – a membraneless germ granule structure which contains components responsible for silencing of transposable elements in the developing germline^42,43^. In addition to Vas, these components include Piwi proteins and their associated small Piwi-interacting RNAs (piRNAs), and Tudor (Tud) domain-containing proteins. To determine whether the anterior nurse cells assembled higher amount of the whole nuage, and not just that of Nv-Vas, we generated antibodies against the *Nasonia* orthologs of Piwi/Aub (Nv-Aub) and the multi-Tud domain scaffold Tud protein (Nv-Tud). Both antibodies were confirmed to specifically react with the target proteins in extracts from ovaries and early embryos, and were validated in RNAi knockdown experiments (Extended Data Fig. 4).

In *Drosophila*, Aub and Tud uniformly colocalize with Vas to nuage in all nurse cells and the posterior germ plasm of the oocyte and early embryos. In contrast to this, we used super-resolution microscopy to show that all these nuage components (Vas, Aub and Tud) are assembled at higher amounts in the nuage around the nuclei of anterior nurse cells in the wasp and the levels of all these proteins are decreased in the posterior nurse cells (Fig. 3a). These data support the mechanism which leads to a higher-level assembly of the whole nuage in *Nasonia*, not just its individual components, in the anterior compartment.

**Fig. 3.**
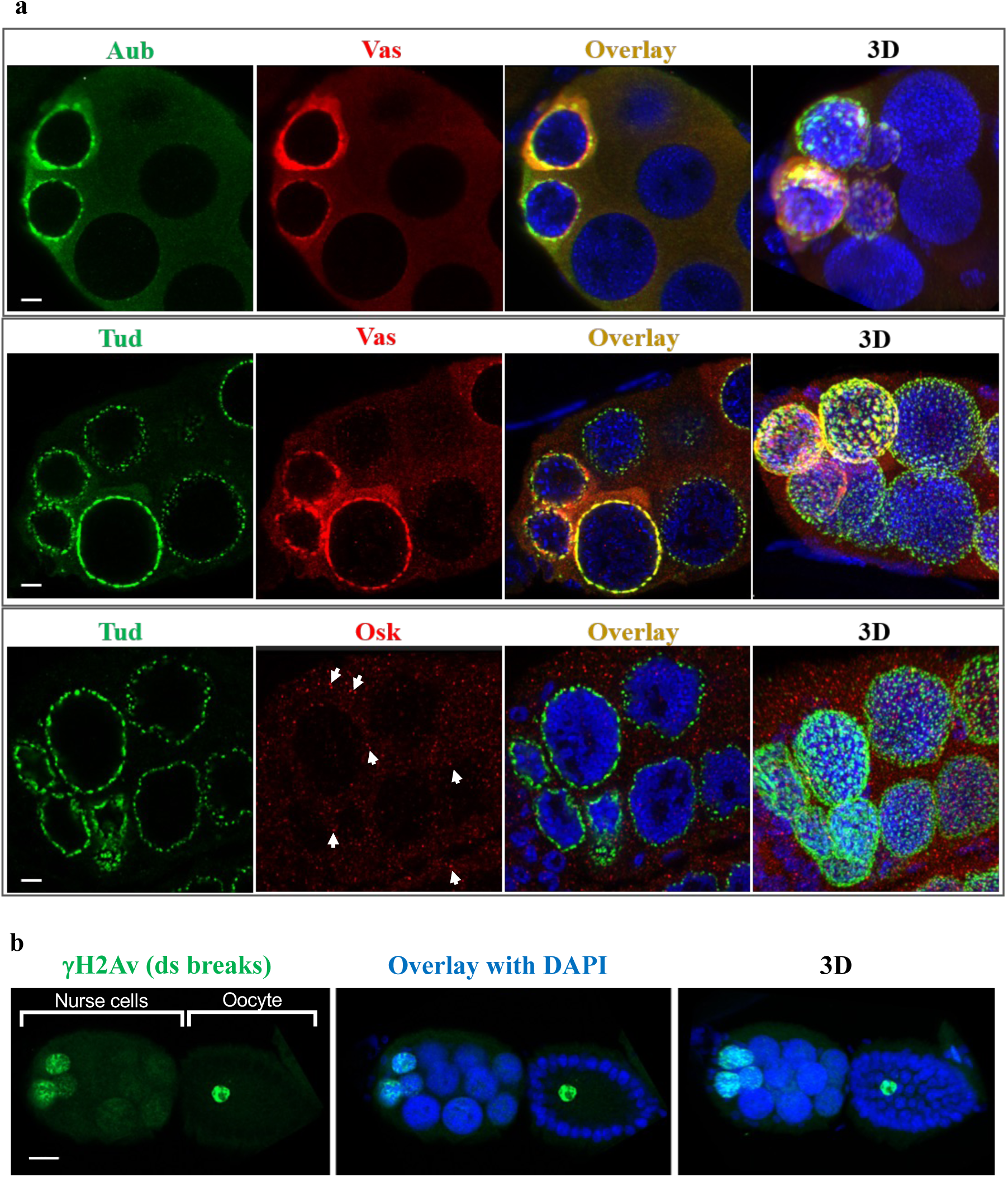
Nurse cells in *Nasonia* egg chambers are different in respect to nuage levels and DNA damage. **a**, Super-resolution microscopy imaging of Long Nv-Vas, Nv-Aub and Nv-Tud in nurse cell compartment shows a high-level assembly of all these components into nuage organelle in anterior nurse cells and substantially lower nuage amounts in posterior nurse cells. 3D images are composite images from multiple optical sections. Also, contrary to *Drosophila*, Nv-Osk is expressed in nurse cells and is assembled into cytoplasmic granules ∼200 nm in diameter (low panel, red channel, indicated with arrowheads). **b**, Anterior nurse cells are stained with anti-yHistone H2Av antibody (green), which is a marker of DNA double-strand breaks, with no significant staining detected in posterior nurse cells. Also, oocyte nucleus shows anti-yHistone H2Av staining. In all images, anterior is to the left. Scale bar in **a** is 5 μm and in **b** it is 15 μm.

We then asked what might be a reason for a substantially high level of nuage assembly in anterior nurse cells in the wasp. Since mobilization of transposable elements can be associated with DNA double-strand breaks (DSBs)^44–46^, we first immunostained *Nasonia* ovaries with anti-γHistone H2Av antibody, which is a marker of the double-strand breaks^47^. Interestingly, we detected a strong γH2Av staining of anterior nurse cells’ nuclei (Fig. 3b, Extended Data Fig. 5a), which show increase from early to later stages of oogenesis, indicating that the same population of nurse cells that assemble high amounts of nuage, shows high level of DSBs.

Given this result, we hypothesized that the reason for a high-level nuage assembly in anterior might be to provide an extra protection of anterior nurse cells from integration of transposable elements into genome which may be needed since high occurrence of DSBs may enhance this integration or potentially lead to activation of transposable elements^48–50^. Therefore, we tested if transposable elements are still silenced in anterior nurse cells despite a high accumulation of DSBs. In support of our hypothesis, in RNA *in situ* experiments detecting expression of a transposon-related gene in *Nasonia* ovaries, we found no evidence that transposable elements are selectively upregulated in anterior nurse cells (Extended Data Fig. 5b), suggesting that high assembly of anterior nuage is needed to effectively silence transposable elements despite the prevalence of DSBs in anterior.

In *Drosophila* nurse cells and early oocyte, *osk* RNA is translationally repressed until its translation is activated at posterior pole of the oocyte at stage 9 of oogenesis and this repression is crucial for normal embryo patterning^51,52^. However, we found that Nv-Osk protein is produced in the cytoplasm of nurse cells and there it is assembled in granules of ∼200 nm in diameter (Fig. 3a). In addition to the lack of detectable interaction between Nv-Osk and Nv-Vas, this Nv-Osk protein expression in nurse cells suggests a different strategy used by *Nasonia* to assemble germ granules and ensure egg and embryonic development.

#### Oosome architecture changes during development from a structure with the uniform distribution of its protein components in oocyte’s posterior to the organelle with Tudor shell and internal Vasa/Aubergine/Oskar granules in preblastoderm embryos

Availability of new antibodies generated in this study against *Nasonia* specific Long Vas isoform, Osk, Tud and Aub, made it possible to analyze the assembly and distribution of these proteins in the oosome at different stages from egg, to the embryonic stage of primordial germ cell formation and provide insights into how this unusually large organelle forms and delivers its components to primordial germ cells when they form at the embryo’s posterior. In the early oosome assembled at the posterior pole of the developing oocyte, all these proteins show generally uniform distribution within the oosome and show a high degree of colocalization (Pearson’s correlation coefficients range from 0.85 to 0.98, Figs. 4a, c). This early oosome is tightly anchored to the posterior cortex of the oocyte and exhibit internal protein free channels or cavities (Fig. 4a). However, after the egg is laid, the oosome is no longer connected to the posterior cortex and starts migrating in the posterior half of the preblastoderm embryo (Fig. 1g, 4b). In stark contrast with the oocyte oosome, in these early embryos, the oosome proteins change their distribution – Nv-Tud is concentrated at the periphery of the oosome, forming a fibrillar shell while Nv-Osk, Long Nv-Vas and Nv-Aub coalesce inside the oosome to form distinct spherical granules ∼400 nm in diameter (Figs. 4b, d, Extended Data Fig. 6). The spherical shape of these granules indicates that they are liquid-like condensates forming in the oosome. In the granules, Nv-Osk, Nv-Vas and Nv-Aub show a substantial colocalization, which is defined as the location of the proteins within the same granules (Figs. 4b, 5b, Extended Data Fig. 6c).

**Fig. 4.**
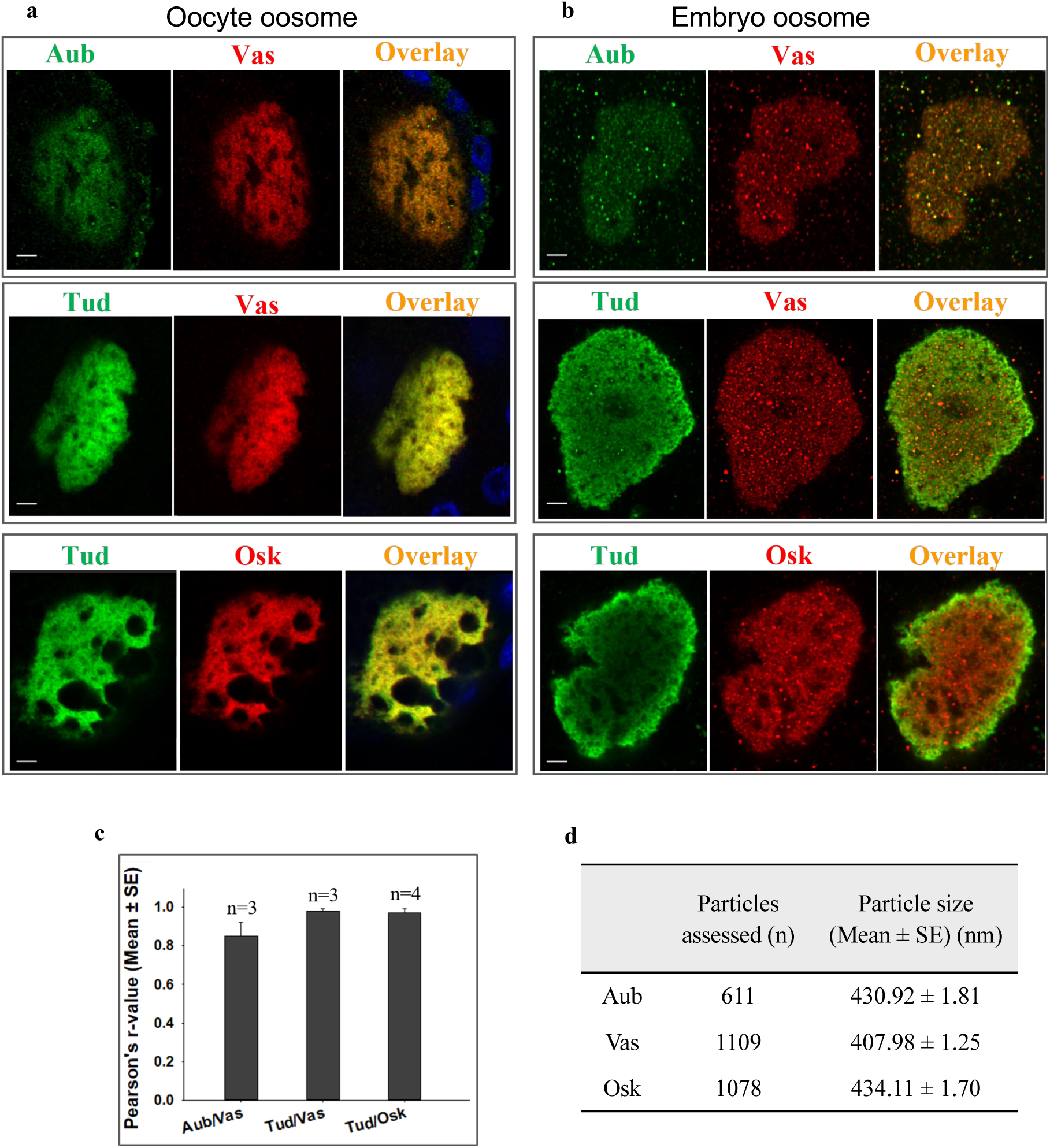
Oosome changes during germline development from a structure with the uniform distribution of its protein components in oocyte to the organelle with Tudor shell and internal Vasa/Aubergine/Oskar granules in preblastoderm embryos. (**a, b**), Super-resolution microscopy imaging of Nv-Aub, Long Nv-Vas, Nv-Tud, and Nv-Osk in the oosome from oocyte’s posterior (**a**) and early preblastoderm embryos (**b**). Overlay images in (**a**) also show DAPI nuclear staining of neighboring follicle cells. While early oosome shows a uniform distribution of its protein components in the oocyte (**a**), in early embryos, the oosome assembles Tud shell and internal Vas/Aub/Osk granules (**b**, Extended Data Fig. 6). Scale bars are 3 μm. **c**, Pearson’s correlation coefficients (mean ± s.e.m.) show high level of colocalization of indicated proteins in the oocyte’s oosome (0.85 ± 0.07, 0.98 ± 0.01 and 0.97 ± 0.02 for Aub/Vas, Tud/Vas and Tud/Osk respectively). Several oosomes, imaged with super-resolution methodology, were quantified for each pair of proteins (n). **d**, Calculation of the size of the Aub, Long Vas, and Osk granules assembled in the oosome’s core in early (preblastoderm) embryos. Diameter of the granules (“particle size”) were measured from super-resolution microscopy images with Imaris software. Number of granules used for calculation is indicated (n).

**Fig. 5.**
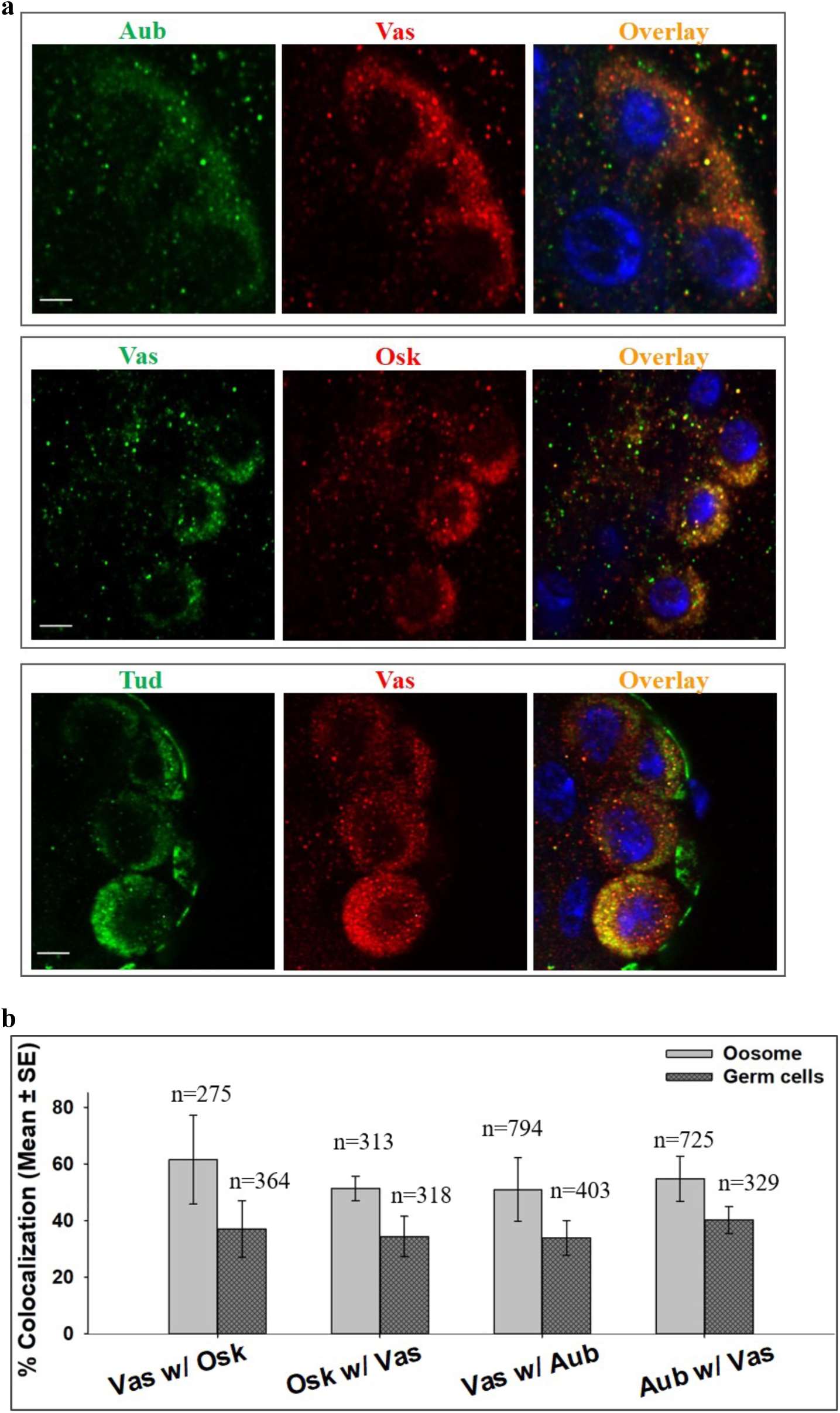
Aubergine, Vasa, Tudor and Oskar-containing granules segregate from the oosome to the cytoplasm of primordial germ cells in *Nasonia*. **a**, Super-resolution microscopy images of immunostained primordial germ cells formed at the posterior tip of the embryo, which show multiple cytoplasmic granules inherited from the oosome containing Nv-Aub, Nv-Vas, Nv-Tud and Nv-Osk as indicated. In Vas/Osk stainings, rabbit anti-Nv-Vas antibody was used and in the other stainings, shown in this Figure, guinea pig anti-Nv-Vas antibody (Long Nv-Vas-specific) was used. Overlay images include DAPI nuclear stain (blue). Scale bars are 3 μm. **b**, Y-axis shows percentages of granules with a given protein that also contain the second protein in the same granules as indicated (mean ± s.e.m.). These colocalization values are calculated for the oosome’s core granules at preblastoderm embryos and for the granules in the cytoplasm of germ cells. Statistical analysis indicated that the differences in the colocalization values for the oosome’s core granules and germ cells’ granules are not significant (p-values for Nv-Vas granules that also show Nv-Osk (Vas w/Osk), Nv-Osk granules that also show Nv-Vas (Osk w/Vas), Nv-Vas granules that also show Nv-Aub (Vas w/Aub) and Nv-Aub granules that also show Nv-Vas (Aub w/Vas) were 0.25, 0.26, 0.28 and 0.21 respectively (Supplementary Table 1). Numbers of particles quantified for colocalization analysis with Imaris software, are indicated for each condition (n).

#### *Nasonia* Aubergine, Vasa, Tudor and Oskar granules segregate from the oosome to the cytoplasm of primordial germ cells

When primordial germ cells form at the embryo’s posterior, Nv-Tud is assembled into granules, which are similar to Nv-Vas, Nv-Aub and Nv-Osk granules formed in the oosome core, and no longer shows the enrichment in the periphery seen in the early embryonic oosome (Fig. 5a). Similarly, Nv-Vas, Nv-Aub and Nv-Osk granules are segregated to the cytoplasm of the germ cells from the oosome (Fig. 5a) and, while it appears that these proteins are less colocalized in germ cells (present in the same granule) than in the oosome before germ cell formation stage (Fig. 5b), this difference is not statistically significant (p-values ranging from 0.21 to 0.28).

#### Disassembly of Tudor shell when embryonic nuclei migrate through the oosome’s core supports the dense shell/liquid core architecture of the oosome

To gain further insights into the organization of the oosome, we took advantage of a specific embryonic developmental stage immediately before germ cell formation. During this stage, embryonic nuclei are migrating to the cortex of the embryo and the nuclei that reach the posterior pole are eventually incorporated into the primordial germ cells. In addition to the nuclei, the oosome components must segregate into the germ cells. Using super-resolution microscopy imaging of the specific stage, just before germ cell formation, we found that the Tud shell breaks down and the nuclei enter the core of the oosome (Fig 6a). In addition to the visualization of this important stage, the nuclei entering the oosome serve as internal probes of the oosome density, which is consistent with overall liquid core, through which the nuclei are able to move, and a more dense Tud shell, which may be flexible, but still needs to disintegrate before nuclei can penetrate the oosome.

**Fig. 6.**
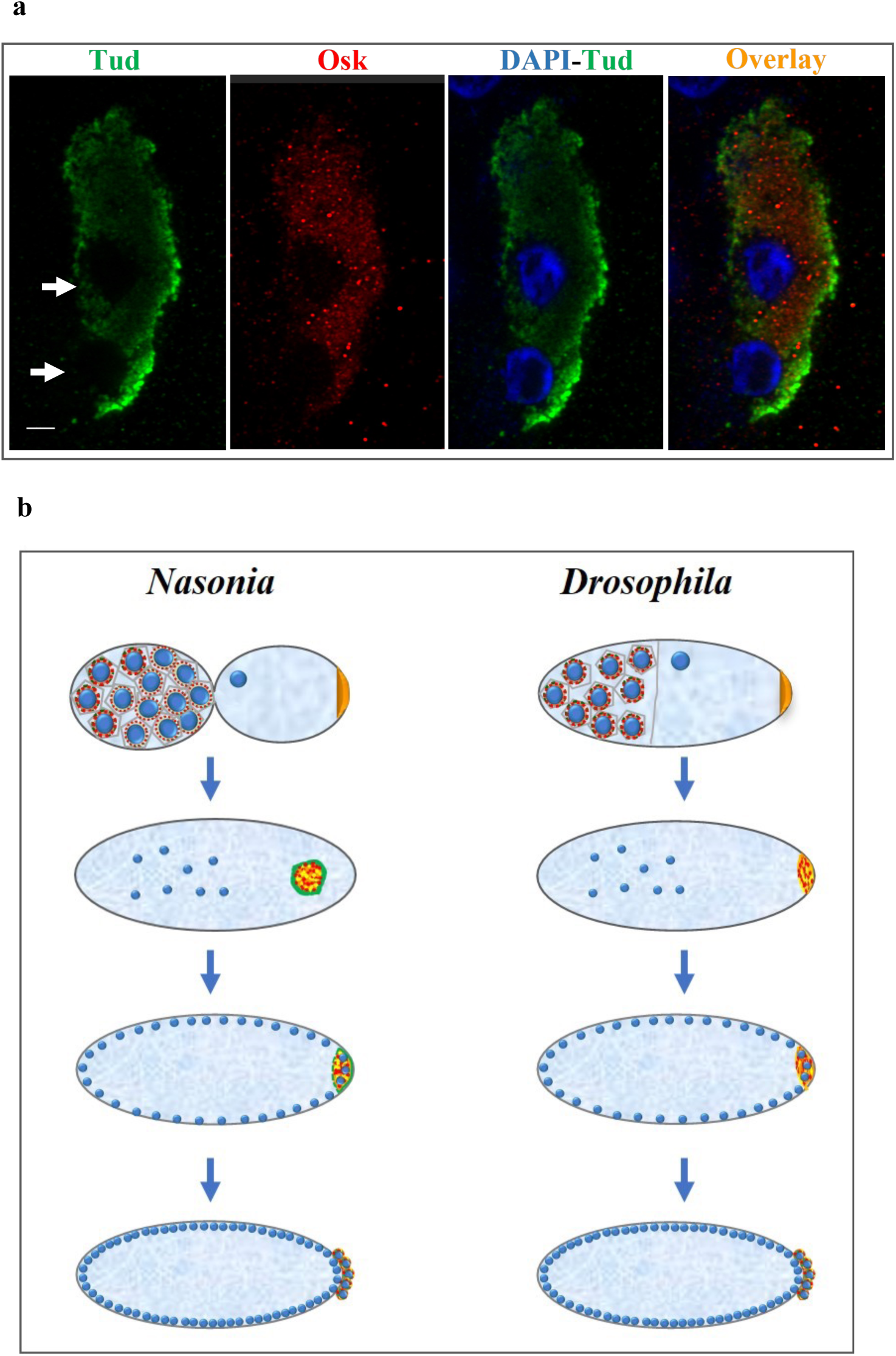
Tudor shell of the oosome breaks down when nuclei migrate through the oosome’s core prior to germ cell formation, consistent with the dense shell/liquid core architecture of the oosome. **a**, Posterior pole of *Nasonia* embryo imaged with super-resolution microscopy just before germ cell formation. Nv-Tud shell (green) breaks down (arrows) at the points of entry of two nuclei (DAPI, blue) which would give rise to primordial germ cells. Oosome core with Nv-Osk granules (red) accommodates the nuclei and spherical shapes of the granules indicate their liquid-like properties in the oosome core. Scale bar is 3 μm. **b**, A diagram of nuage and the oosome assembly steps in *Nasonia* compared with nuage and polar granules in *Drosophila*. Top images show egg chambers with higher nuage accumulation in *Nasonia* anterior nurse cells compared to posterior nurse cells while in *Drosophila*, nuage is equally assembled in nurse cells (nuage and nuclei are indicated in red and blue respectively). The rest of the images show embryonic stages with the oosome forming Tud shell (green) and Osk/Vas/Aub core granules (red) whereas similarly sized *Drosophila* polar granules (red) contain all these proteins. Subsequently, in *Nasonia*, the Nv-Tud shell breaks down when nuclei of future germ cells enter the oosome at the posterior pole and Nv-Tud is incorporated together with other proteins into primordial germ cells. In contrast, in *Drosophila*, nuclei of future primordial germ cells enter the germ plasm without its apparent structural change. Also, in *Drosophila*, germ plasm remains to be anchored at the posterior pole during early embryogenesis until germ cell formation.

## Discussion

In this study, we analyzed how a large solitary membraneless granule, the oosome, is assembled in the wasp *Nasonia vitripennis*, which is strikingly different from the multiple small homologous structures formed in germ cells of *Drosophila melanogaster* (Fig. 6b). An intriguing question is how the integrity of the oosome, which is by itself has a size of an individual cell, is maintained without the membrane? The need for a special mechanism to maintain the oosome integrity is especially apparent during the dramatic movement of this organelle in the embryo before it reaches the posterior pole to segregate its components to primordial germ cells – the germline progenitors.

In particular, we showed that the oosome is initially assembled in the oocyte’s posterior where it is associated with the posterior cortex and shows a homogeneous distribution of its several protein components including Tud, Osk, Vas and Aub. However, during early embryogenesis, we found that Tud protein becomes enriched at the periphery of the oosome and there it assembles a fibrous shell, which encapsulates the core content of the oosome.

Using a super-resolution microscopy in the oosome core, we detected spherical granules of about 400 nm in diameter, which contain *Nasonia* homologs of Aub, Vas and Osk. The spherical shape of these granule, point to their liquid characteristics and their formation inside the oosome core via liquid-liquid phase separation mechanism. The size of the granules is similar to the polar granules assembled in *Drosophila* germ plasm, however, the embryonic polar granules are more amorphous and usually not spherical consistent with the observations which indicate that *Drosophila* polar granules may have hydrogel-like phases^40^.

We demonstrate that the assembly of Tud shell coincides with the migration of the oosome in the posterior half of the early embryo and that the Tud envelope starts breaking down when nuclei of future germ cells begin to enter the oosome prior to their formation. These data suggest that Tud protein shell fulfills the role of a membrane in this organelle, and helps to maintain a unique molecular environment in the oosome liquid core and the integrity of the whole oosome. Also, these data show that similarly to *Drosophila* polar granules, the core of the oosome contains multiple similarly sized granules which contain Aub, Vas and Osk, which however, in contrast to *Drosophila* germ plasm, are assembled in a specialized Tud protein bound compartment (Fig. 6b).

Protein envelopes are assembled in different systems in nature most notably in viral capsids or to protect a viral DNA in a host cell or around membrane vesicles during vesicular transport. However, oosome may provide a unique example of a protein shell which assembles around a compartment of ∼20 micrometers in diameter, which far exceeds known protein shells. The surface area of the oosome, which needs to be covered by a Tud shell is about 1300 square microns. If a single molecule particle size for *Drosophila* Tud^53^ (which has a similar molecular mass to *Nasonia* Tud) is used (∼11 nm) then to generate a monolayer with Tud molecules around the oosome core, about 10 million Tud protein molecules would be needed. Future research will need to address the structural details of the Nv-Tud shell and whether Nv-Tud is sufficient to form a shell by itself.

In *Drosophila*, Osk protein is a crucial germ plasm inducer, which directly interacts with Vas in germ plasm, recruits it to the granules in S2 cells and it is necessary and sufficient to assemble germ plasm and form germ cells^17,20,37,40^. Similarly, in *Nasonia*, Osk is required for the oosome formation^18^, however, our data indicate that this protein employs a different mechanism to form the oosome. In particular, while Nv-Osk often colocalizes with Nv-Vas in the oosome and primordial germ cells, in S2 cells, Nv-Osk assembles granules by itself, and does not recruit Nv-Vas when the proteins are coexpressed. Consistent with the inability of Nv-Osk to recruit Nv-Vas to its granules formed in heterogenous S2 cell system, we could not co-immunoprecipitate Nv-Osk and Nv-Vas from *Nasonia* ovarian extracts. While these data are consistent with lack of direct interaction between Nv-Osk and Nv-Vas, we cannot rule that interaction out, and more experimentation will be needed to determine if Nv-Osk and Nv-Vas directly interact in the oosome or they colocalize in the core granules because of indirect interactions mediated by RNA or other proteins.

Development of *Nasonia* egg chambers show several unusual molecular and cellular features in comparison to those of *Drosophila*. First, we demonstrated that anterior nurse cells in *Nasonia* are different from their posterior counterparts as they assemble higher amounts of perinuclear nuage (Fig. 6b). Interestingly, we found that the anterior nurse cells’ nuclei also showed a higher number of foci labeled with anti-yHistone H2Av antibody, which is a marker of the DSBs. The occurrence of DSBs in a distinct population of nurse cells in any organisms has not been reported before to our knowledge, and our data suggest that higher nuage assembly in those cells may be needed to provide extra protection against transposable elements which would, otherwise, mobilize more efficiently in the presence of damaged DNA.

Furthermore, in contrast to *Drosophila*, we found that *Nasonia* ovaries express two alternatively spliced Vas isoforms, which are different from each other by 96 amino-acid segment enriched in FG repeats. Our circular dichroism measurements of the purified 96 amino-acid fragment, show a CD spectrum which is typical for a disordered protein, indicating that the Long Vas isoform contains an intrinsically disordered region, which, similarly to FG-rich nucleoporins, may engage in the formation of mesh-like structures in perinuclear nuage and the oosome by engaging aromatic groups of phenylalanines in intermolecular *π*-*π* interactions similar to those in the nuclear pore complex. Consistent with this mechanism, using a Long Vas-specific antibody, we demonstrated that Long Vas is efficiently assembled in nuage and the oosome core. Moreover, while both Long and Short Vas isoforms are expressed equally well in ovaries, the Long isoform became the major Vas isoform in early embryos indicating that the oosome mainly contain the Long Vas.

In conclusion, the assembly of the oosome, an unusually large and complex membraneless organelle, relies on the combination of highly conserved components, such as Nv-Osk, Nv-Vas, Nv-Aub, and Nv-Tud on one hand, as well as a suite of novel features, including a novel Nv-Vas isoform, an unusual shell of Nv-Tud protein demarcating the edges of the oosome, and unusual distribution of nuage in the ovary. Understanding how these features work together to produce the unusual features of the oosome, and help it to carry out its function in establishing the germline will provide important insights into how evolution works at the subcellular level to bring about changes in conserved cellular processes.

## Methods

### Protein expression and purification

To express *N. vitripennis* proteins, different constructs were generated using *N. vitripennis* embryo cDNA encoding Nv*-*Vas full-length protein (amino acids 1-865), Nv*-*Vas Long isoform-specific region (amino acids 19-114), Nv-Osk LOTUS domain (amino acids 16-90), Nv-Aub (amino acids 695-893), and Nv*-*Tud (amino acids 2104-2431). Each construct was cloned separately into the pET SUMO vector using the TA cloning method (Thermo Fisher Scientific). The resulting plasmids were verified by DNA sequencing.

The plasmids with respective constructs were transformed into *E. coli* BL21 (DE3) competent cells (New England Biolabs, C2527), which were incubated on Luria Bertani (LB) agar plates containing kanamycin (50 μg/ml). The primary culture of the transformed single bacterial colony containing specific plasmid was cultured overnight at 37 °C with vigorous shaking at 230 rpm in the LB broth medium containing kanamycin. The secondary cultures were grown by adding 1% of primary cultures with kanamycin at 37 °C with shaking at 250 rpm.

When the bacterial density (OD600) reached 0.60, the secondary culture was allowed to cool down for 30 min before inducing protein expression with Isopropyl β-D-1-thiogalactopyranoside (IPTG) (0.5 mM). The culture was then incubated overnight at 16 °C. After the overnight expression of recombinant proteins, the cultures were centrifuged at 8,000 g at 4 °C for 15 min, the growth media was discarded and the harvested cells were saved at −20 °C until further experiments.

The protein purifications of Nv-Vas Long isoform-specific fragment, Nv-Osk LOTUS domain, Nv-Aub, and Nv-Tud were carried out essentially as described^26^ with slight modifications. In particular, the harvested cells were resuspended in lysis buffer (50 mM Tris/Tris-HCl, 500 mM NaCl, 5% glycerol, 0.1 mg/ml lysozyme, 1% Triton X-100, protease inhibitors with 1mM phenylmethanesulfonyl fluoride (PMSF), pH 7.4) and incubated at 4 °C for 1 hr for lysis. The lysate was sonicated on ice using an ultrasonicator for 3 min (10 sec on/off), followed by centrifugation of the lysate at 14,000 g, 4 °C for 15 min. The pellets were discarded and the supernatant was filtered and loaded onto the Ni-NTA column pre-equilibrated with the column buffer (50 mM Tris/Tris–HCl, 500 mM NaCl, 5% glycerol, pH 7.5) and incubated at 4 °C for 1 hr. The unbound protein was washed with 10-50 mM imidazole solutions in the column buffer. The proteins were eluted with 100 and 200 mM imidazole.

For the purification of Nv*-*Vas full-length protein, the above method was modified since the protein formed inclusion bodies in bacterial cells. Specifically, cells with the recombinant protein were resuspended in lysis buffer and incubated at 4 °C for 1 hr for lysis. The lysate was sonicated on ice using an ultrasonicator for 3 min (10 sec on/off), followed by centrifugation of the lysate at 14,000 g, 4 °C for 15 min. Then the supernatant was discarded and the pellets were again resuspended in buffer with 0.5% SDS in 50 mM Tris/Tris-HCl, 500 mM NaCl, 5% glycerol, 0.1 mg/ml lysozyme, 1% Triton X-100, pH 7.4. Then the filtered protein extract was loaded onto the Ni-NTA column pre-equilibrated with the column buffer and the protein was purified following further purification steps described above.

### Production of antibodies, affinity purification, and validation

The purified proteins in the form of soluble liquid antigens were submitted to Cocalico Biologicals, Inc. Pennsylvania, USA for antibody production. The antibodies against full-length Nv-Vas, Nv-Aub, and Nv-Tud were raised in rabbits while those against Nv-Vas Long isoform-specific region and Nv-Osk LOTUS domain were raised in guinea pigs. The standard protocol for antibody raising in both animals included initial inoculation of the antigen on the first day (0 d), the first boost after 14^th^ day, the second boost on 22^nd^ day, the first test antisera obtained on 35^th^ day, the third boost on 49^th^ day, the second antisera obtained on 56^th^ day, the final antisera obtained on 91^st^ day.

The antisera from 56^th^ day or 91^st^ day were further affinity purified against purified proteins with glutathione S-transferase (GST) affinity tags. The 26 kD GST tag was fused with the N-terminus of the recombinant protein. The purified recombinant protein was immobilized onto glutathione-agarose using a crosslinker disuccinimidyl suberate (DSS) which contains an amine-reactive N-hydroxysuccinimide (NHS) ester at each end of an 8-carbon spacer arm. Then the crude serum was passed through the crosslinked beads, washed with a non-amine-containing buffer (100 mM sodium phosphate, 150 mM NaCl (pH 7.2), and then the purified antibody was eluted using low pH elution buffer (0.2 M glycine HCl, pH 2.6). Antibody purification protocol was adopted and modified from Zalazar et al (2014)^54^ and Thermo Fisher Scientific DSS user manual.

The antibodies were validated with western blot methodology against wild-type control and RNAi knockdown embryos or ovaries. The methodology for generating RNAi knockdowns in the wasp has been described^55^.

### Circular Dichroism analysis of Long Vasa-specific FG-rich region

To purify a tagless Nv-Long Vas-specific FG-rich region (aa 19-114) for CD analysis, after the His-tag purification, the His-SUMO tag was removed by SUMO protease. Subsequently, the tagless protein was purified with ion exchange chromatography with HiTrap DEAE Sepharose Fast Flow anion exchange column (Cytiva) with Akta Pure liquid chromatography system (GE Healthcare) using linear 0-1 M NaCl gradient in 50 mM Tris-HCl (pH 7.75). CD spectrum of the protein (0.28 mg/ml in 50 mM Tris-HCl, pH 7.75) was obtained at 200-260 nm at 20 °C in Jasco quartz cuvette (1 mm pathlength) using Jasco J-1500 CD spectrophotometer.

### *Nasonia oskar* 5′-UTR mapping

To determine whether *Nv*-*osk* 5′-UTR has a previously unknown extended region, which codes for a ∼6 kD extension detected with our Nv-Osk antibody, RT-PCR reactions with ovarian RNA and several forward primers targeting the genomic region upstream of previously predicted *Nv*-*osk* 5′-UTR were carried out followed by sequencing of RT-PCR products to confirm intronless cDNA products. In these reactions, the common reverse primer (5′-GCCGCCCTTTCTTGTCAGTA-3′), that anneals to the exon 2 of *Nv*-*osk* RNA, was used. The locations of the upstream forward primers, which amplified the cDNA corresponding to the properly spliced *Nv*-*osk* RNA are indicated in Extended Data Fig. 3 and their sequences are as follows. upstr_Nvosk2_fwd (5′-GGTCTTCCCGAAGCTGTGCTTC-3′) and upstr_Nvosk3_fwd (5′-GGAAGGGTGAATCGCGCATG-3′).

### In vitro expression of the *Nasonia* Oskar

An *Nv*-*osk* cDNA with extended new 5′-coding sequence that codes for the additional 57 N-terminal amino acids reported in this work was generated in an RT-PCR reaction using *Nasonia* embryonic RNA. This cDNA was cloned into the pET SUMO vector using the TA cloning method as described above for the generation of other constructs. Subsequently, using the plasmid as the DNA template, a PCR fragment containing the complete *Nv-osk* coding sequence was synthesized for protein expression with PURExpress® In Vitro Protein Synthesis Kit (New England Biolabs) using manufacturer’s instructions. After in vitro translation, ∼51 kD Nv-Osk protein product was detected with anti-Nv-Osk antibody in a western blot experiment.

### Immunoprecipitation of *Nasonia* Oskar and Vasa

IPs with guinea pig anti-Nv-Vas Long isoform and anti-Nv-Osk antibodies, crosslinked to protein A/G magnetic beads with DSS, were performed using ovarian and embryonic lysates in lysis buffer (PBS, with EDTA-free protease inhibitors, 1 mM PMSF, 0.05% IGEPAL CA-630) at 4 °C overnight. Subsequently, the beads were washed and proteins bound to the beads were eluted according to the manufacturer’s instructions for Pierce Crosslink Magnetic IP/Co-IP Kit (Thermo Scientific, cat # 88805) followed by western-blot detection as specified in figure legends.

### *Nasonia* ovary and embryo collection

*Nasonia* colonies were maintained at 25 °C in culture tubes with *Sarcophaga bullata* (grey flesh fly) pupae (Carolina Biological Supply). At 25 °C, the wasp completes its life cycle in 15 days. The ovaries were collected from 2-4-day-old adults. Embryos were collected essentially as described previously^56^. In particular, egg-laying chambers (the waspinators) were made out of petri dishes by making holes in the lid to insert about 1-inch cut pipette tips so that *Sarcophaga* pupae can be held upright inside the tips. Accordingly, the mated 2-4-day-old *Nasonia* females were exposed to the *Sarcophaga* host pupae placed inside the tips and, subsequently, early wasp embryos (0-6 hrs) were collected by crack opening the host pupae. The *Nasonia* ovaries and embryos were collected in PBS and PBS with 0.1% Triton X-100 (PBT) respectively.

### Immunohistochemistry and RNA FISH

For immunostaining, *Nasonia* embryos and ovaries were fixed with 5% formaldehyde (made in PBT) in heptane for 20 min at room temperature (∼22 °C) then overnight at 4 °C. The embryos were manually devitellinized using a sharp needle. The embryos and ovaries were stained with the primary antibody overnight at 4 °C (1:250 for guinea pig anti-Nv-Osk antibody; and 1:1000 for the other antibodies generated in this work), then with Alexa Fluor 488-or Cyanine3 (Cy3)-labeled secondary antibodies (1:500) for 2 hours at room temperature. For detection of DSBs, mouse anti-yHistone H2Av antibody (DSHB)^47^ (1:200) was used.

RNA FISH experiments were done using Stellaris oligonucleotide probes labeled with Quasar 570 Dye according to the manufacture’s protocol (LGC Biosearch Technologies). Sequences of the probes are listed in Supplementary Table 2.

### Expression of *Nasonia* proteins in *Drosophila* S2 cells

For expression of Nv-Osk and Nv-Vas proteins in S2 (*Drosophila* Schneider 2) cells (Expression Systems, cat # 94-005S), corresponding cDNAs were subcloned under control of Actin5C promoter (GenScript). The resulting constructs had N-terminal Halo or EGFP tags (Nv-Osk) and mCherry (Long and Short Nv-Vas isoforms). Transfection of the S2 cells was done as described previously^40^ and for HaloTag-Nv-Osk imaging, Janelia Fluor HaloTag Ligand 646 (Promega) was used.

### Super-resolution imaging

Super-resolution imaging was performed with Zeiss LSM 880/Airyscan super-resolution module system and inverted laser scanning confocal microscope AxioObserver and Plan-Apochromat ×63/1.4 Oil DIC M27 objective at Vanderbilt University Cell Imaging Shared Resource (CISR). Images were acquired as a Z-stack and analyzed with Imaris software (version 9.5, Oxford Instruments) and HP Z8 workstation. To measure the size of the spherical granules assembled by a given protein in the oosome, automatic 3D reconstructions of the granules were performed using Spots tool in Imaris. Colocalization of the proteins’ distribution in the oocyte’s oosome was quantified with method of Costes^57^ utilizing Coloc 2 module of ImageJ 1.53t, NIH. Percentage of co-occurrence of proteins in the same granules in the embryo was quantified using Spots colocalization analysis tool of the Imaris software.

### Statistics

For the co-occurrence analysis of proteins in the same granules, the percentage of the granules, containing a given protein pair, was then transformed into angular values^58^ for statistical analysis. The normal distribution of data was examined using *proc univariate* features of the SAS® 9.2 software (SAS Institute Inc. 2013, SAS Institute, Cary, NC). After that, the data were subjected to Student’s t-test (two-sample independent t-test) using the Proc ttest feature of the SAS software to compare the mean values of protein co-occurrence in the same granules between preblastoderm embryo’s oosome and germ cells. The data presented in the text are all untransformed mean co-occurrence percentage values for each protein pair (Fig. 5b, Supplementary Table 1).

## Data availability

The datasets used during the current study are available from the corresponding authors upon reasonable request.

## Acknowledgments

We thank J. Schafer and the Vanderbilt Cell Imaging Shared Resource for help with Zeiss LSM 880/Airyscan super-resolution microscopy. Also, we thank I. Antoshechkin and P. Ferree for their help with identification of a potential transposon-related gene expressed in the *Nasonia* ovary. In addition, we thank Abigail Collins for help with some experiments. This study was supported by NIH National Institute of General Medical Sciences grant (R01GM129153) to J.A.L. and A.L.A. This work was partly supported by a grant from the NIH National Institute of General Medical Sciences, P20GM103436.

## Author contributions

Conceptualization, A.L.A., J.A.L.; Methodology, K.K, S.J.T., A.K., A.L.A., J.A.L.; Experimentation: K.K., S.J.T., A.K., R.S., E.A.; Analysis, A.L.A., J.A.L., K.K., S.J.T., A.K., R.S.; Writing – first draft, A.L.A., K.K., J.A.L.; Writing – review/editing, all authors; Funding acquisition, A.L.A, J.A.L; Supervision, A.L.A., J.A.L.

---

**Extended Data Fig. 1.**
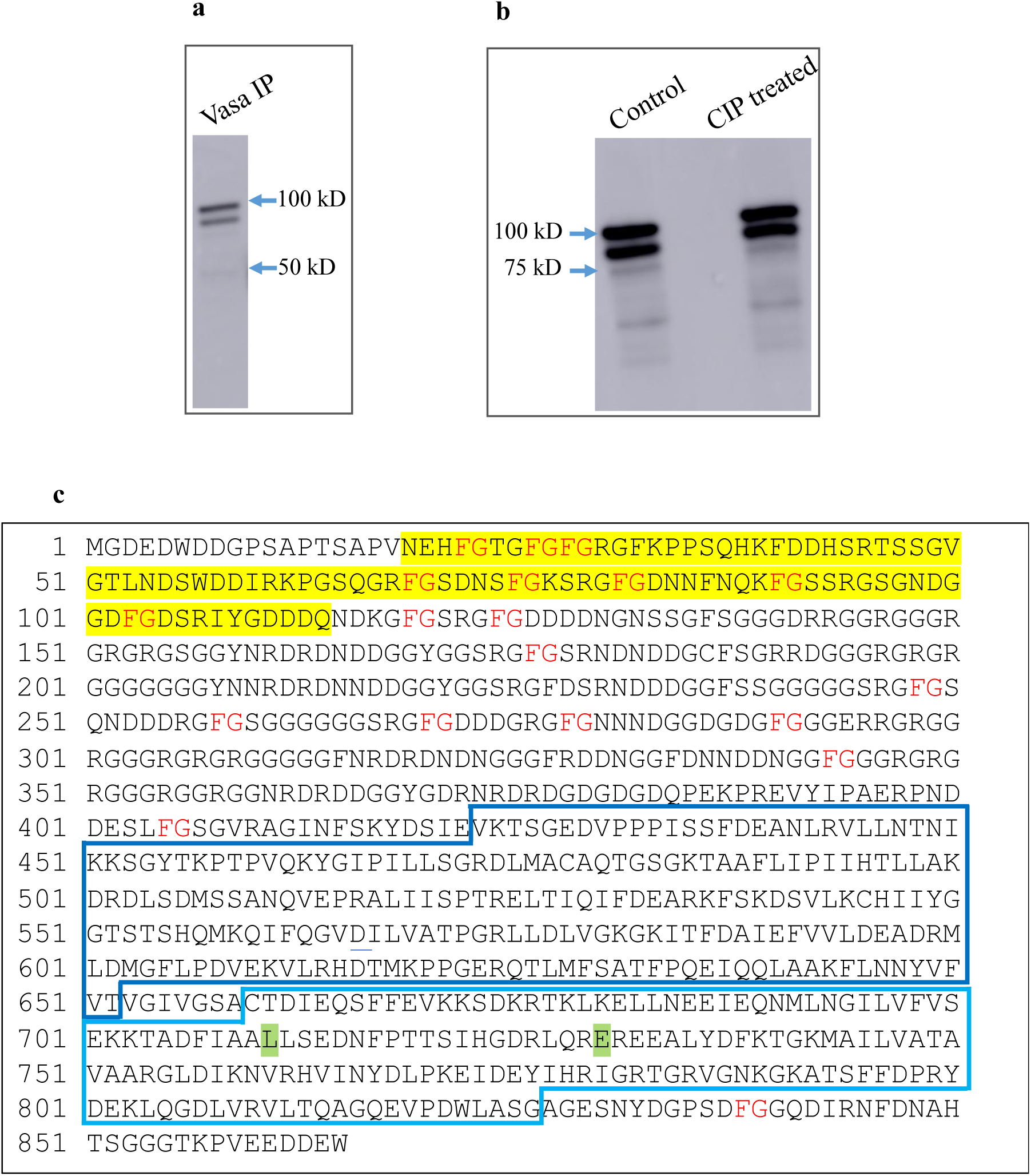
**a**, IP with rabbit anti-Nv-Vas antibody, resulted in the isolation of both Nv-Vas isoforms (Nv-Vas identity of both protein bands was confirmed by mass spectrometry). **b**, Western-blot images after detection with rabbit anti-Nv-Vas antibody show no difference in the relative levels of Nv-Vas isoforms after *Nasonia* ovarian lysate was treated with CIP enzyme as compared with control (ovarian lysate treated with CIP buffer only). This indicates that different isoforms are not generated by phosphorylation of a single Nv-Vas polypeptide. **c**, Sequence of Long Nv-Vas isoform with Phe-Gly (FG)-rich region, absent in Short Nv-Vas isoform, highlighted in yellow. FG repeats are indicated along the entire sequence. The region absent in Short Nv-Vas, contributes 11% of the total length of Long Nv-Vas, however, it contains 42% of all the FG motifs of the Long Nv-Vas. The locations of two conserved helicase RecA-like domains are indicated with boxes. Also, consistent with our finding of no detectable association between Nv-Vas and Nv-Osk proteins, some amino acids shown to be important for interaction of *Drosophila* Vas with Osk or for *Drosophila* Vas localization to germ granules, have not been conserved in Nv-Vas. Specifically, Phe508 in *Drosophila* Vas, which is important for interaction with Osk^38^, is replaced with Leu in the corresponding position 711 of Nv-Vas. Similarly, Gln527 in *Drosophila* Vas, which is required for Vas recruitment to germ granules in *Drosophila*^41^, corresponds to Glu at position 730 of Nv-Vas. Both Leu711 and Glu730 are located in RecA-like C-terminal domain and are highlighted in green. These molecular differences, between *Drosophila* and *Nasonia* Vas proteins, in addition to the FG repeats, which are absent in *Drosophila* Vas, may point to a distinct mechanism of Nv-Vas recruitment to the oosome which differs from the *Drosophila* Vas assembly into germ granules.

**Extended Data Fig. 2.**
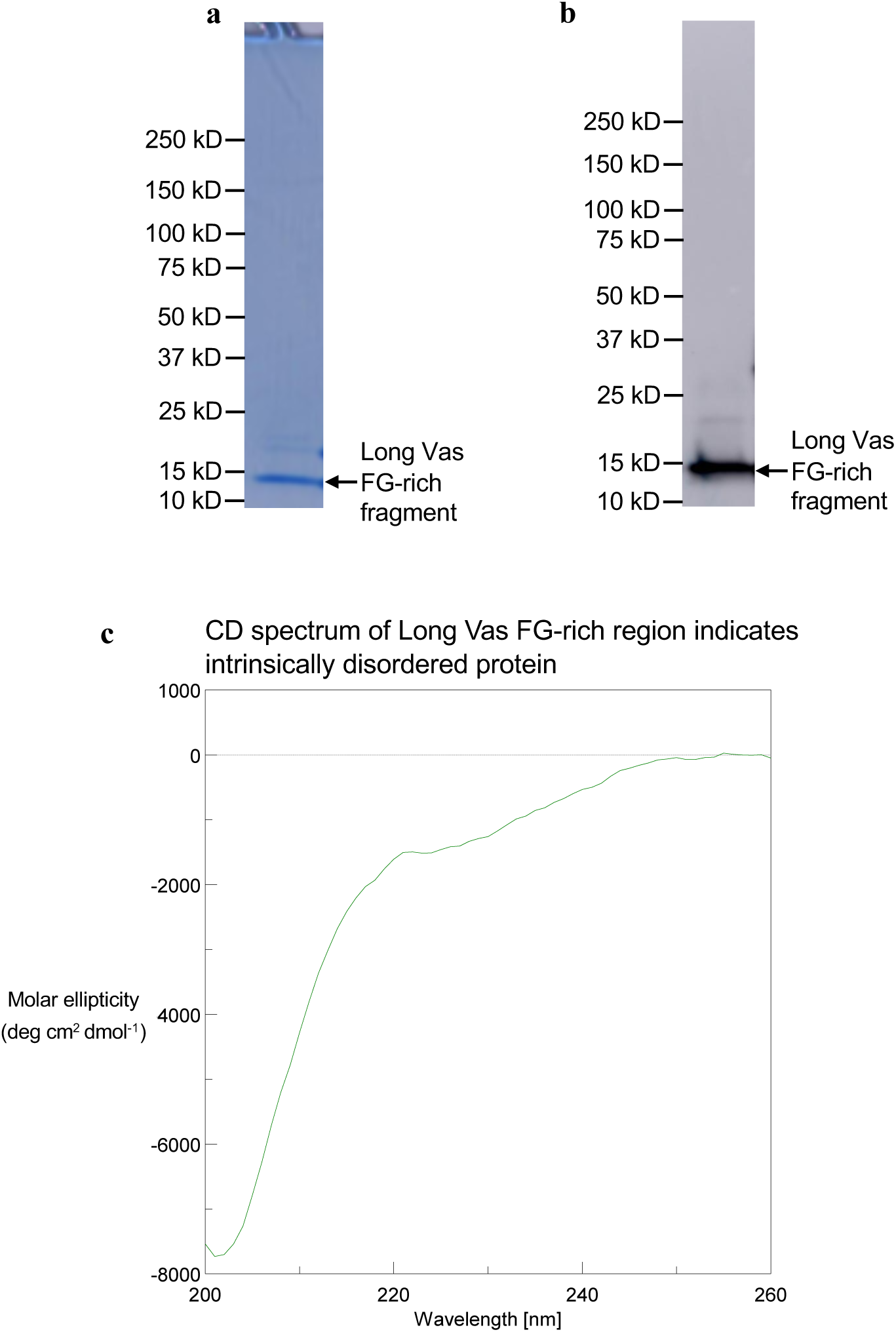
Circular dichroism analysis of purified Long Nv-Vas-specific FG-rich region indicates that this is an intrinsically disordered segment. **a**, Coomassie-stained polyacrylamide gel shows purified tagless Long Nv-Vas-specific FG-rich fragment. **b**, Western-blot experiment of purified protein shown in (**a**), using anti-Long Nv-Vas-specific FG-rich region antibody confirms the identity of purified fragment. **c**, CD spectrum of purified Long Nv-Vas-specific FG-rich fragment (0.28 mg/ml in 50 mM Tris-HCl, pH 7.75) is characteristic for unfolded proteins and closely mirrors CD spectra of disordered FG-rich regions of nucleoporins^27,28^. These spectra show large minimum close to 200 nm and low ellipticity around 230 nm.

**Extended Data Fig. 3.**
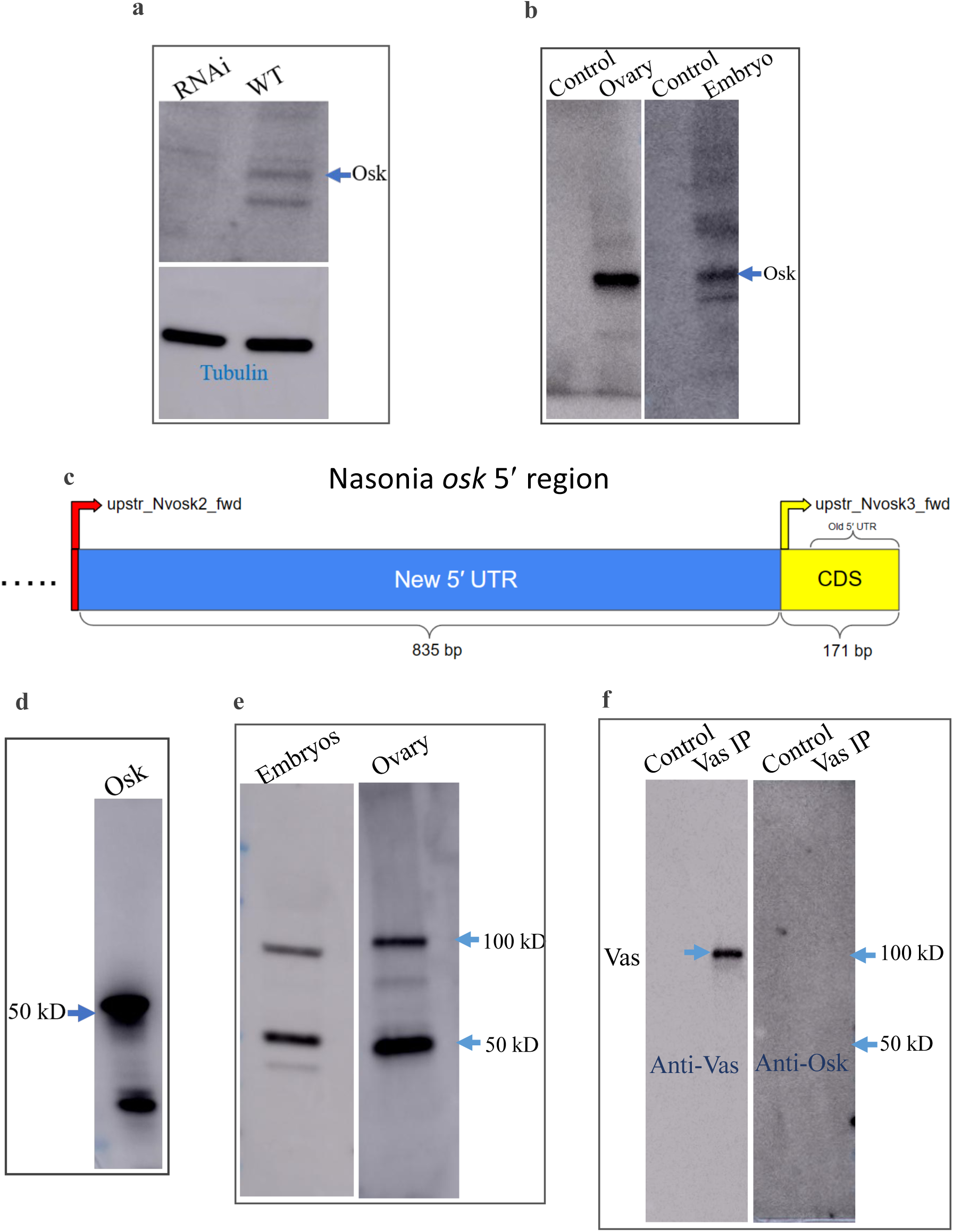
Molecular analysis of *Nv*-*osk* 5′ region and Nv-Osk protein. **a**, Top: western-blot image shows reduction of ∼51 kD Nv-Osk in *Nv*-*osk* RNAi embryonic knockdown samples validating new anti-Nv-Osk antibody. Bottom: loading control with anti-β-Tubulin antibody demonstrates equal loading of RNAi and wild-type embryonic extracts during western-blot experiment. **b**, Western-blot detection of ∼51 kD Nv-Osk after immunoprecipitation with anti-Nv-Osk antibody from both wasp ovarian and embryonic lysates. Control lanes correspond to experiments performed with empty beads (without anti-Nv-Osk antibody) under the same conditions as the IP experiments. **c**, A schematic of *Nv*-*osk* 5′ UTR mapping experiments. To probe a potentially extended 5′ UTR of *Nv*-*osk* mRNA, several forward primers were used in separate RT-PCR reactions using RNA purified from ovaries and a downstream reverse primer annealed to exon 2 of *Nv*-*osk* RNA. Locations of two forward primers, which generated cDNA products corresponding to intronless *Nv*-*osk* RNA, as verified by sequencing, are indicated on the *Nv*-*osk* 5′ gene region with arrows. The new coding sequence (CDS), which codes for additional 57 N-terminal amino acids of Nv-Osk (NTD) is shown in yellow. This new CDS region, which includes previously reported 5′ UTR is indicated. **d**, Detection of ∼51 kD Nv-Osk, generated with bacterial in vitro translation system, by anti-Nv-Osk antibody in a western-blot experiment; **e**, In addition to ∼51 kD Nv-Osk, in some experiments, less intense bands around 100 kD are detected in both ovarian and embryonic samples, consistent with Nv-Osk dimerization. **f**, Nv-Osk does not co-IP with Nv-Vas. Nv-Vas IP was conducted by crosslinking anti-Long Nv-Vas-specific antibody with magnetic protein A/G beads with DSS crosslinker. Subsequently, the ovarian extract was incubated with the crosslinked beads and bound proteins were probed with anti-Long Nv-Vas and anti-Nv-Osk antibodies as shown in left and right panels respectively. The same procedure was followed for the control sample except empty beads (without the antibody) were used.

**Extended Data Fig. 4.**
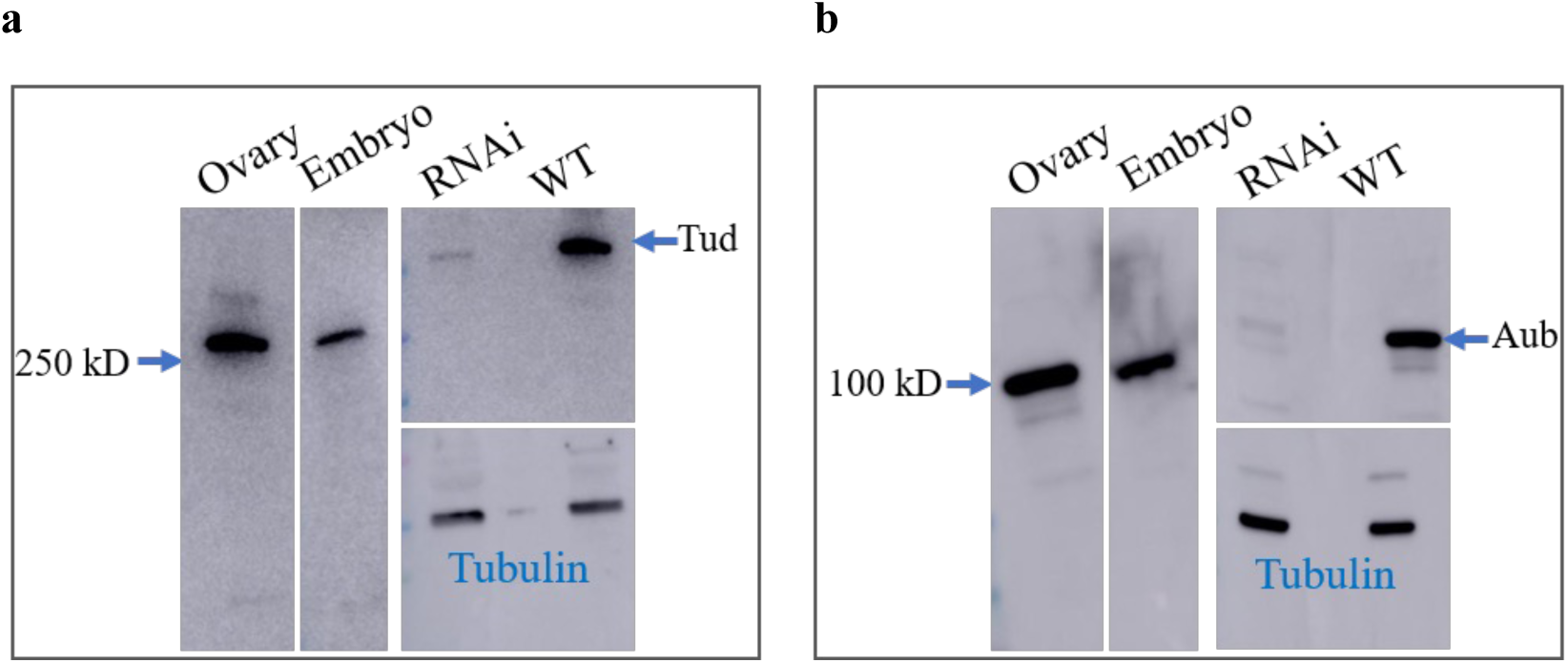
Validation of anti-Nv-Tudor and anti-Nv-Aubergine antibodies. **a**, Western-blot detection of 275 kD Nv-Tud (UniProt ID A0A7M7QCB7) in ovaries (left panel) and embryos (middle panel) with rabbit anti-Nv-Tud antibody. Reduction of Nv-Tud in the *Nv*-*tud* RNAi ovarian sample compared to wild type control, detected with the antibody, confirms that the antibody recognizes Nv-Tud (top right). Loading controls are done with anti-β-Tubulin antibody (bottom right). **b**, Western-blot detection of 102 kD Nv-Aub (UniProt ID A0A7M7G3W4) in ovaries (left) and embryos (middle) with rabbit anti-Nv-Aub antibody. Reduction of Nv-Aub in the *Nv*-*aub* RNAi ovarian sample compared to wild type control is detected with the antibody confirming its specificity (top right). Loading controls are performed with anti-β-Tubulin antibody (bottom right).

**Extended Data Fig. 5.**
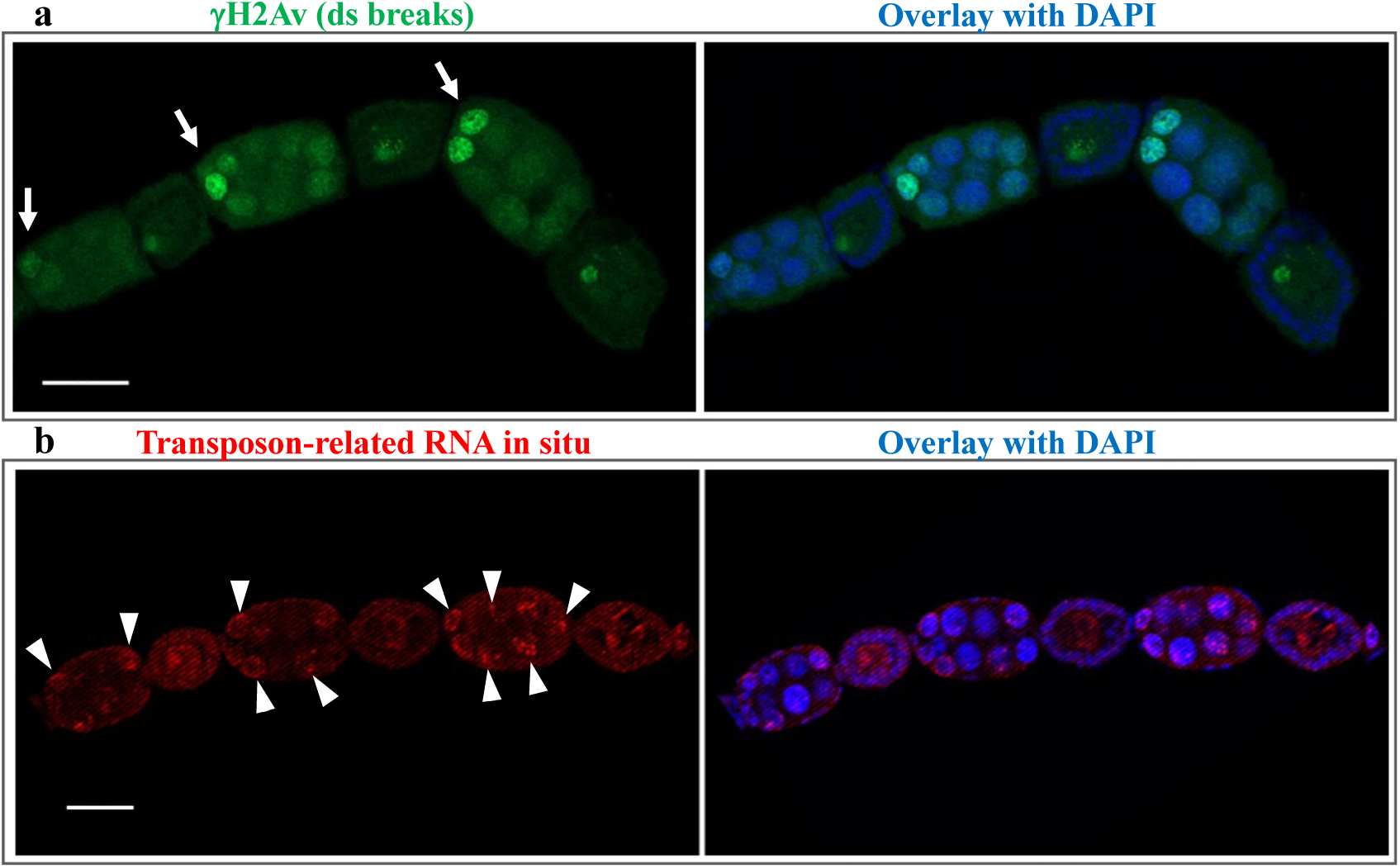
Accumulation of double-strand DNA breaks in anterior nurse cells does not appear to lead to overexpression of transposable elements. **a**, *Nasonia* egg chambers stained with anti-yHistone H2Av antibody (green), a marker of DNA double-strand breaks, show accumulation of double-strand breaks in anterior nurse cells during oogenesis (indicated with arrows). **b**, RNA FISH experiments detect low uniform expression of a transposable element-related gene (LOC103317460) in all nurse cells (red, arrowheads), which was used based on the sequence match to transposon-related RNA, referred to as NV30926, previously found to be expressed in *Nasonia* ovaries^61^. Sequences of Stellaris FISH oligonucleotide probes labeled with Quasar 570 Dye, which were used in this experiment, are listed in Supplementary Table 2. Scale bars in **a** and **b** are 40 μm.

**Extended Data Fig. 6.**
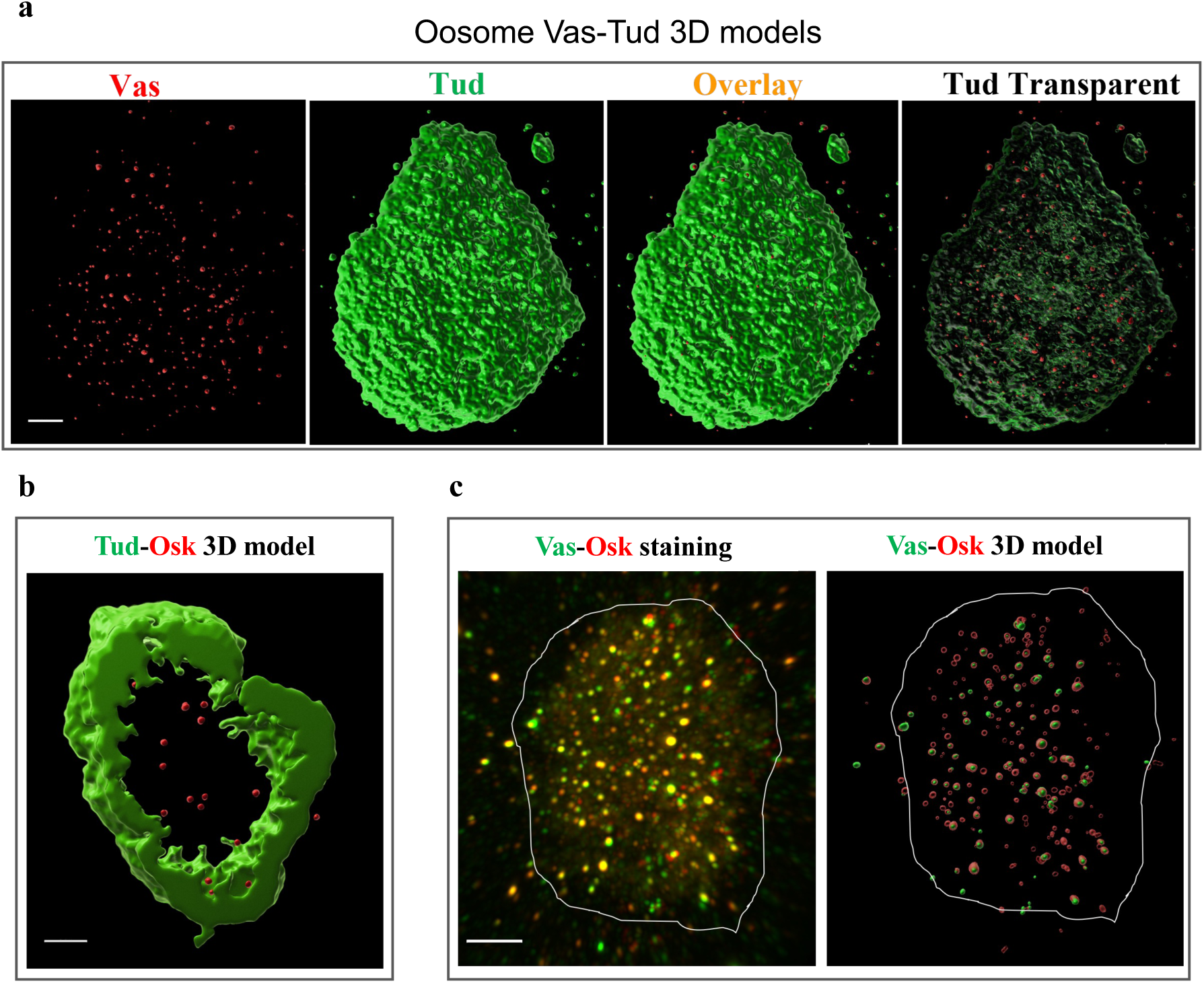
3D reconstruction of a Tudor shell and Vasa/Oskar core granules of the oosome. **a**, 3D model of Nv-Tud and Long Nv-Vas distribution in preblastoderm embryos was generated from multiple super-resolution microscopy optical sections with Imaris software. Nv-Vas forms core granules (red) enveloped by Nv-Tud shell (green). In overlay image, Long Nv-Vas core granules are inside the solid Nv-Tud cage, and become visible when the Nv-Tud shell is made transparent. Scale bar is 5μm. **b**, 3D reconstruction of the internal segment of the embryo’s oosome based on super-resolution optical sections from Nv-Tud/Nv-Osk immunostaining experiments. Nv-Tud shell and internal core Nv-Osk granules can be seen. **c**, Left: super-resolution microscopy image of preblastoderm embryo’s oosome stained with rabbit anti-Nv-Vas (green) and guinea pig anti-Nv-Osk (red). While rabbit anti-Nv-Vas antibody recognizes both Long and Short Nv-Vas isoforms, in the embryos, predominantly long Nv-Vas isoform is expressed (Fig. 1a). Right: 3D modeling based on the optical sections from the left panel. White outline in the images indicates approximately the border of the oosome. In (**b**, **c**) scale bars are 3 μm.

**Supplementary Table 1.** Statistical analysis of co-occurrence of proteins in the same granules in the oosome and primordial germ cells of Nv embryo.

| Proteins | % granules (Mean ± SE) | | t-test | *p-value* |
| --- | --- | --- | --- | --- |
|  | Oosome | Germ cell |  |  |
| Osk w/Vas | 51.28 ± 4.36 | 34.39 ± 7.16 | -1.26 | 0.26 |
| Vas w/Osk | 61.42 ± 15.64 | 37.01 ± 10.04 | -1.28 | 0.25 |
| Aub w/Vas | 54.73 ± 7.98 | 40.22 ± 4.75 | -1.42 | 0.21 |
| Vas w/Aub | 50.94 ± 11.28 | 33.90 ± 6.16 | -1.19 | 0.28 |

**Supplementary Table 2.**
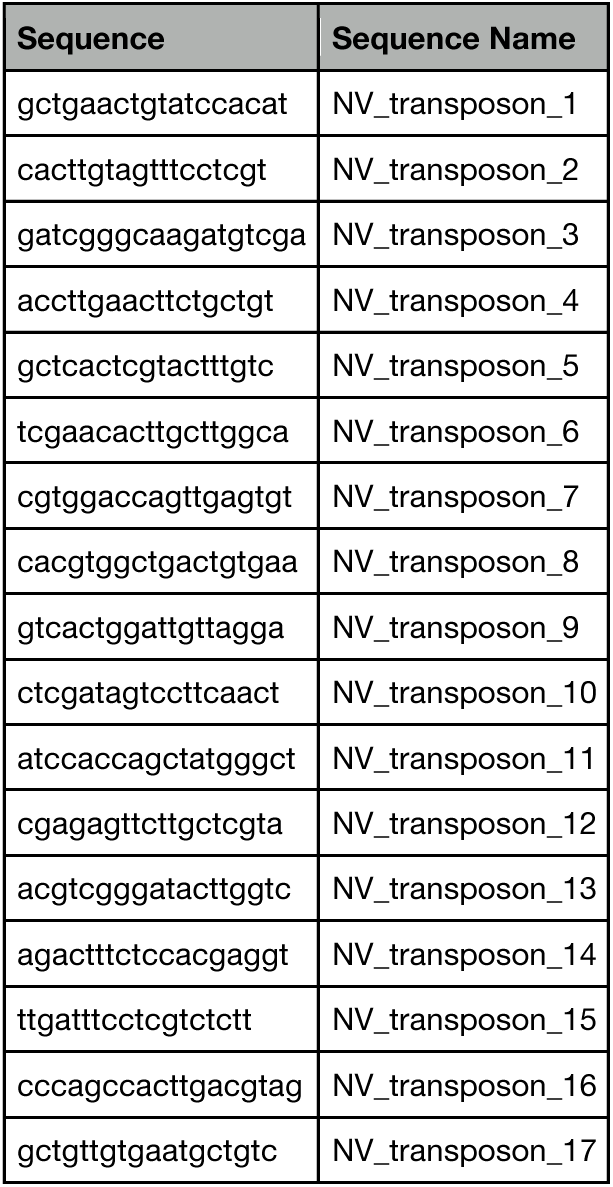
Stellaris oligonucleotide probes used in RNA FISH experiments.

